# S100A4 Is a Targetable Regulator of Immune Suppression in Glioblastoma

**DOI:** 10.64898/2026.11.10.765528

**Authors:** Thomas L Wong, Jia-Shiun Leu, Maryam Faisal, Nourhan Abdelfattah, Wei Xiong, Fransica Leonard, Yaqoob Ali, Dante Jimenez, Han Nhat Tran, Sung Yun Jung, Sinem Gunalp, Reece Kang, Jose Maldonado, Zhiqiang An, Ningyan Zhang, Kyuson Yun

## Abstract

Glioblastoma (GBM) is a highly aggressive brain tumor that remains largely refractory to immunotherapy owing to a profoundly immunosuppressive tumor microenvironment (TME). Here, we identify S100A4 as an actionable regulator of immune suppression and demonstrate that its inhibition converts GBM into an immunotherapy-responsive state. Genetic knockdown of *S100a4* in glioma cells reprogrammed the GBM TME, enhancing CD8⁺ T cell infiltration and effector function while promoting maturation of antigen-presenting cells. These changes translated into significant survival benefits across diverse molecular subtypes of GBM. Therapeutically, a novel S100A4/transferrin receptor (TFR) bispecific antibody phenocopied *S100a4*-KD and showed significant therapeutic efficacy, particularly in combination with PD-1 blockade. Notably, BsA+anti-PD1 combination treatment generated tertiary lymphoid-like structures in GBM. Together, these findings establish S100A4 as a targetable driver of immune evasion in GBM and provide a translational strategy for overcoming resistance to immunotherapy in this lethal disease.

## INTRODUCTION

Glioblastoma (GBM) remains the deadliest adult primary brain tumor with high risk of recurrence and very limited treatment options (*1*). While immunotherapies have shown significant clinical benefit in other cancers, they have largely failed to demonstrate benefit for human GBM patients in multiple clinical trials (*2–5*). Nivolumab (αPD-1) failed to improve overall survival (OS) or progression-free survival (PFS) in newly diagnosed GBM when compared against or combined with GBM standard of care (*2, 3*). It also failed to improve survival in recurrent GBM compared to bevacizumab (*5*). Resistance to immune checkpoint inhibitors (ICIs) is largely attributed to an immunosuppressive GBM tumor microenvironment (TME) that is predominantly occupied by suppressive myeloid cells (*6–9*) and the paucity of cytotoxic T cells necessary for ICI response(*10, 11*). Other challenges to treating GBM include: a) intrinsic tumor heterogeneity that influences therapy response (*7, 12*), b) physiological barriers such as the blood-brain barrier (BBB) that restrict treatments, including antibody drugs, from reaching the tumor tissue (*13, 14*), and c) the immunosuppressive nature of GBM standard-of-care, which includes lymphocyte-depleting radio-chemotherapy and high-dose immunosuppressive corticosteroids (*15–19*). Thus, the most promising immunotherapeutic strategies currently in clinical evaluation are either delivered locally to directly reach tumor tissues, such as oncolytic virotherapy(*20–22*), or act systematically to overcome immunosuppression and promote anti-tumor immunity, such as neoantigen-mRNA/DNA/peptide vaccines(*23–25*), or combinatorial immunotherapies (*26, 27*). Therefore, improving immunotherapy efficacy in GBM will require treatments that simultaneously overcome physiological barriers to delivery, such as the BBB, and convert the GBM TME from “immune cold” to “immune hot” tumors.

Tertiary lymphoid structures (TLSs) are ectopic lymphocyte aggregates formed in inflamed tissue regions that resemble secondary lymphoid organs(*28–30*). They organize antigen-presenting dendritic cells, B cells, and T cells to drive T and B cell responses (*28, 29*). In the context of cancer immunity, they may be formed peri- or intra-tumorally, and their presence has been positively associated with patient survival and response to ICI in cancers such as non-small cell lung cancer, breast cancer, melanoma, and renal cell carcinoma(*31–33*). TLSs of varying maturation states are present in adult gliomas, including glioblastoma (*30, 34–36*). While their clinical role in GBM has not been fully determined, their presence is significantly correlated with OS in glioma patients (*34–37*). In CT-2A and GL261 preclinical mouse GBM models, TLS formation is associated with a transition to an immune activated state, enhanced T cell infiltration, and response to ICI(*37, 38*).

We have previously identified S100A4, a multifunctional small Ca^2+^ binding protein of the S100 protein family, as a biomarker and regulator of glioma stem cells and a potential immunotherapy target in GBM(*7, 39–41*). S100A4 expression is induced in response to stress, and it can function both intra- and extra-cellularly, depending on its binding partners. Intracellularly, S100A4 regulates diverse oncogenic processes including invasion, stemness, angiogenesis, and p53 function. Extracellularly, S100A4 can act as a cytokine or damage-associated molecular pattern (DAMP) molecule, binding to cell surface receptors such as RAGE, TLR4, IL10R, and GP130 to activate downstream signaling pathways (*41–44*). In GBM, S100A4 is highly expressed in both cancer cells and immune cells, and higher *S100A4* expression in human GBM tissues predicts worse survival(*7*). We previously showed that deletion of *S100a4* in stromal cells (*S100a4*^-/-^ host mice) is sufficient to significantly extend survival in an aggressive syngeneic mouse glioma model(*7*). On the other hand, *S100a4*-knockdown (*S100a4*-KD) in human and mouse glioma cells reduced stemness and tumor initiating capacity *in vivo* (*7, 39*). Given the emerging studies reporting immune-modulatory function of cancer stem cells (*45–51*), and our studies demonstrating the role of S100A4 in GBM immune suppression in stromal cells, we sought to elucidate the TME modulatory role of S100A4 and evaluate a novel approach to inhibit S100A4 function in GBM.

Here, we report that S100A4 in glioma cells promotes immune suppressive TME through regulation of chemokine/cytokine expression. *S100a4* knockdown reprograms the GBM TME, increasing antigen presentation and CD8 T cell infiltration, sensitizing GBMs to ICI treatment. Additionally, we generated a novel αS100A4/αTfR bispecific antibody (BsA) that utilizes Transferrin-receptor(TfR)-mediated transcytosis to deliver the BsA across the BBB (*52, 53*). The αS100A4/αTfR BsA reprograms the GBM TME and synergizes with ICI to significantly extend survival in multiple GBM subtype mouse models. Notably both *S100a4*-KD and BsA+anti-PD1 treated gliomas increase the density of vasculature with HEV and TLS-like structures in multiple glioma models. Given the challenges associated with immunotherapies in GBM, these results strongly suggest that the αS100A4/αTfR BsA is a highly promising novel immunotherapy for GBM.

## RESULTS

### S100A4 expression elevates at glioma recurrence

S100A4 is highly expressed in human glioma, in both IDH-wild type (IDH-wt) and IDH-mutant (IDH-mut) gliomas, and its high expression is predictive of worse survival(*7*). To assess whether S100A4 expression is associated with glioma recurrence, we analyzed the GLASS consortium dataset, a large longitudinal human glioma dataset with matched primary and recurrent bulk RNA sequencing samples from the same patients (*54–56*). Consistent with a previous report (*7*), we observed that S100A4 expression significantly stratifies patient survival in all-glioma and IDH-mut-glioma cohorts, while there was a trend towards improved survival with lower S100A4 expression in the IDH-wt/GBM glioma cohort (**Figures 1A-C**). We further segregated IDH-mut glioma patients based on S100A4 expression at primary diagnosis or at recurrence (**Figures 1D, E**). We observed that while high S100A4 expression at primary diagnosis shows a trend towards worse survival, this difference became significant at recurrence in IDH-mut glioma (**Figures 1D, E)**. To determine whether S100A4 expression is associated with recurrent glioma, we compared S100A4 expression at recurrence vs primary diagnosis in the same patients. IDH-wt glioma patients overall had higher levels of S100A4 expression compared to IDH-mut gliomas (**Figure 1F**). Additionally, while there was not a significant change in S100A4 expression between matched primary and recurrent samples in IDH-wt glioma patients, for IDH-mut gliomas, with and without codeletion of chromosome 1p19q, S100A4 expression increased at recurrence (**Figure 1F**). By applying multivariable cox proportional hazards model, we confirmed that high S100A4 expression was a negative predictor for overall survival in IDH-wt gliomas and recurrent IDH-mut gliomas independent of age, sex, and other clinical variables (**Supplementary Figure 1A**). Using multivariable logistic regression, we determined that S100A4 expression in all measured cohorts was independent of age and sex, and in the all-glioma cohort was positively associated with IDH-mut 1p19q-non-codeletion and IDH-wt status (vs IDH-mut 1p19q-codeletion) (**Supplementary Figure 1B**).

**Figure 1:**
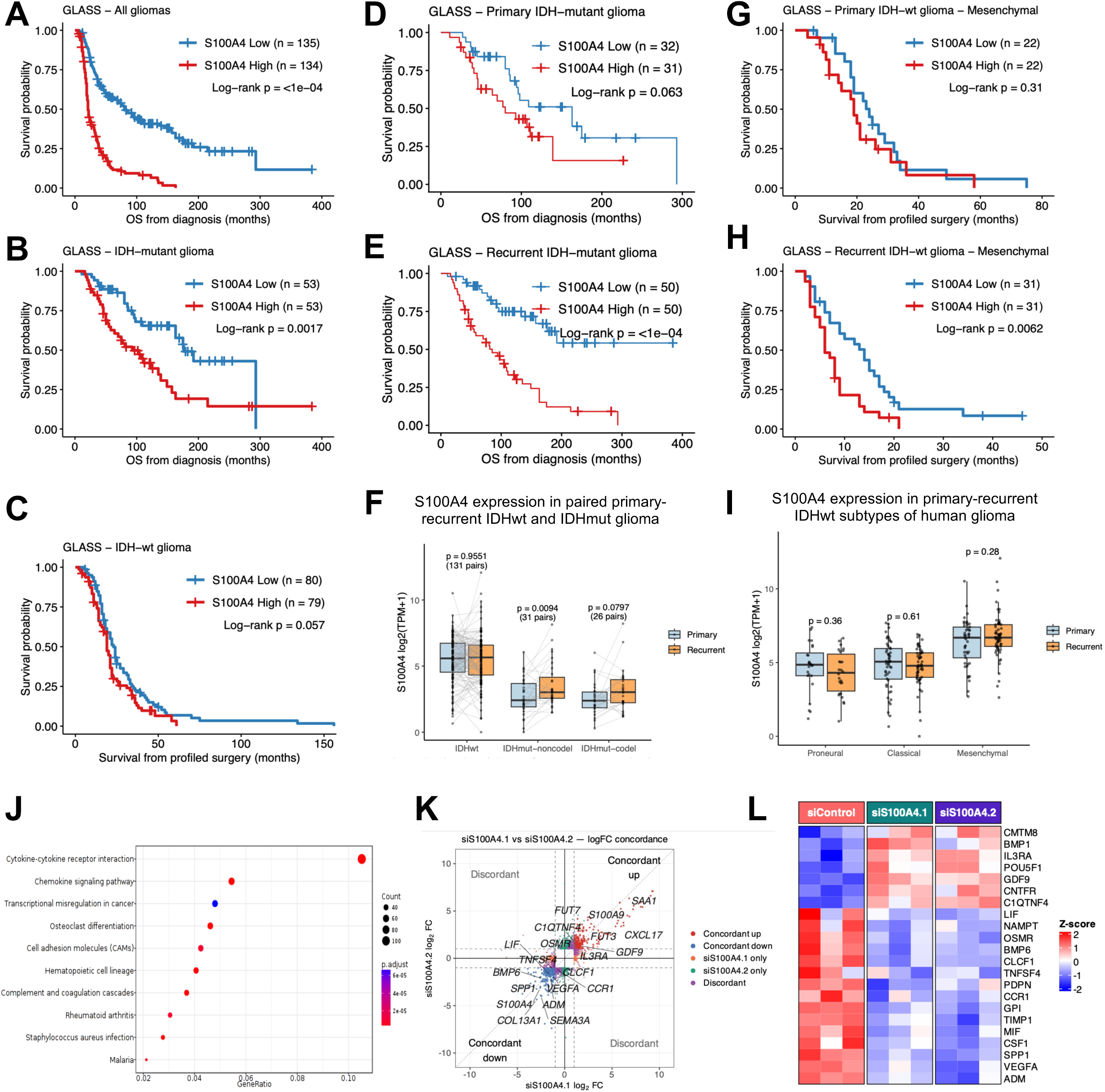
S100A4 expression in human glioma predicts survival and regulates the secretome to modulate the GBM TME. A-E. Kaplan-Meier plot depicting overall survival distributions from diagnosis or profiled surgery of (A) Glioma, (B) IDH-mutant Glioma, (C) IDH-wildtype Glioma, (D) Primary IDH-mutant Glioma, (E) Recurrent IDH-mutant Glioma stratified by high or low S100A4 expression, split determined by median value within cohort. *p*-values determined by log-rank Mantel-cox test. **F.** Paired boxplot depicting S100A4 expression within primary and recurrent IDH-wildtype, IDH-mutant-non-1p19q-codeletion, and IDH-mutant-1p19q-codeletion gliomas. *p-*values determined by paired Wilcoxon signed-rank test. **G-H.** Kaplan-Meier plot depicting overall survival distributions from profiled surgery of (G) primary and (H) recurrent IDH-WT Glioma Mesenchymal stratified by high or low S100A4 expression, split determined by median value within cohort. *p*-values determined by log-rank Mantel-cox test. **I.** Box plots depicting S100A4 expression within primary and recurrent IDH-wildtype gliomas by transcriptional subtype. *p-*values determined by unpaired Wilcoxon signed-rank test. **J.** Over-enrichment analysis showing up regulated GO_BP gene sets for *S100a4* knockdown MMC1 gliomaspheres vs controls. **K.** Scatter plot showing common up and downregulated genes between siS100A4.1 and siS100A4.2 vs controls (n=3 per group). **L.** Heatmap showing normalized expression of selected up and downregulated genes with siS100A4 treatment (n=3 per group).

Additionally, we compared S100A4 expression in IDH-wt/GBM samples by molecular subtypes. In the mesenchymal subtype, the most aggressive and treatment-resistant subtype of human GBM, S100A4 expression significantly stratified patient survival at recurrence but not at primary diagnosis (**Figures 1G-H**). Multivariable cox proportional hazards analysis confirmed that high S100A4 expression was an independent negative predictor for overall survival in this cohort (**Supplementary Figure 1A**). This was not the case in pro-neural and classical subtypes (**Supplementary Figure 1C-F**). Among the GBM subtypes, S100A4 expression was highest in the mesenchymal subtype, but there was no significant difference between primary and recurrent samples for all molecular subtypes (**Figures 1I**). Together, these results suggest that S100A4 expression is strongly associated with the mesenchymal subtype, which is commonly associated with glioma recurrence and high immune infiltration, consistent with our previous report that S100A4 is a master regulator of mesenchymal transition(*39*).

### S100A4 in glioma cells regulates cytokine and chemokine expression and modulates the GBM TME

To elucidate the mechanism of action for S100A4 in glioma cells, we performed siRNA-knockdown (KD) of S100A4 in human primary gliomaspheres (MMC1) and performed differential gene expression (DGE) analysis (**Supplementary Figure 2A**). The most differentially enriched pathway in S100A4-KD cells was “cytokine-cytokine interaction”, suggesting that S100A4-expressing glioma cells produce cytokines and modulate immune environment in GBM (**Figure 1J**). Indeed, top DEGs in *S100A4*-KD include chemokines and cytokines associated with angiogenesis and immune infiltration and activation, including *CSF1, SPP1, VEGFA, POSTN, ADM, TNFSF4/OX40L, and MIF* (**Figures 1K, L**), suggesting that a major function of S100A4 in glioma cells is to modulate the GBM TME by regulating chemokine and cytokine expression.

To test whether S100A4-kD is sufficient to alter immune infiltrates in GBM, we knocked down *S100a4* in mouse gliomaspheres derived from *S100β-vErbB; p53^-/-^* (SVP) mouse model of GBM(*40*). First, we confirmed that S100a4 regulates cytokines in mouse gliomas by analyzing conditioned media from sh-*Ctrl* and sh-*S100a4* gliomaspheres using a cytokine array. *S100a4*-KD decreased SPP1, VEGF, and CCL2 levels in the conditioned media (**Supplementary Figures 2B, C)**. Together, these and our published results indicate that S100A4 function in glioma cells is conserved and that it may promote immune suppressive TME in GBM.

To test this hypothesis *in vivo*, we intracranially injected sh-*Ctrl* and sh-*S100a4 SVP gliomaspheres from* two independent spontaneous gliomas (5459 & 6228) into the striatum of syngeneic C57BL/6 (B6) hosts (**Figure 2A**). Flow cytometry analysis showed that in both 5459 and 6228 tumors, *S100a4*-KD led to an increase in T cell infiltration, with a preferential increase in cytotoxic CD8 T cells (**Figures 2B, C, Supplementary Figure 3**). We further validated this via immunofluorescence (IF) analysis, spatially confirming an increased CD3 and CD8 T cell infiltration in sh-*S100a4* tumors vs controls (**Figure 2D, E)**. Next we evaluated whether *S100a4*-KD affected overall survival in glioma-bearing mice and found that *S100a4* knockdown significantly increased survival in both 5459 and 6228 models (5459: median OS sh-*S100a4* 25 days vs sh-*Ctrl* 21 days, 19% increase in median OS, *p*-value=0.0226; 6228: median OS sh-*S100a4* 34 days vs sh-*Ctrl* 29.5 days, 15% increase in median OS, *p*-value=0.0108) (**Figures 2F, G)**. These results suggest that *S100a4*-KD in glioma cells is sufficient to reprogram the TME and extend survival.

**Figure 2:**
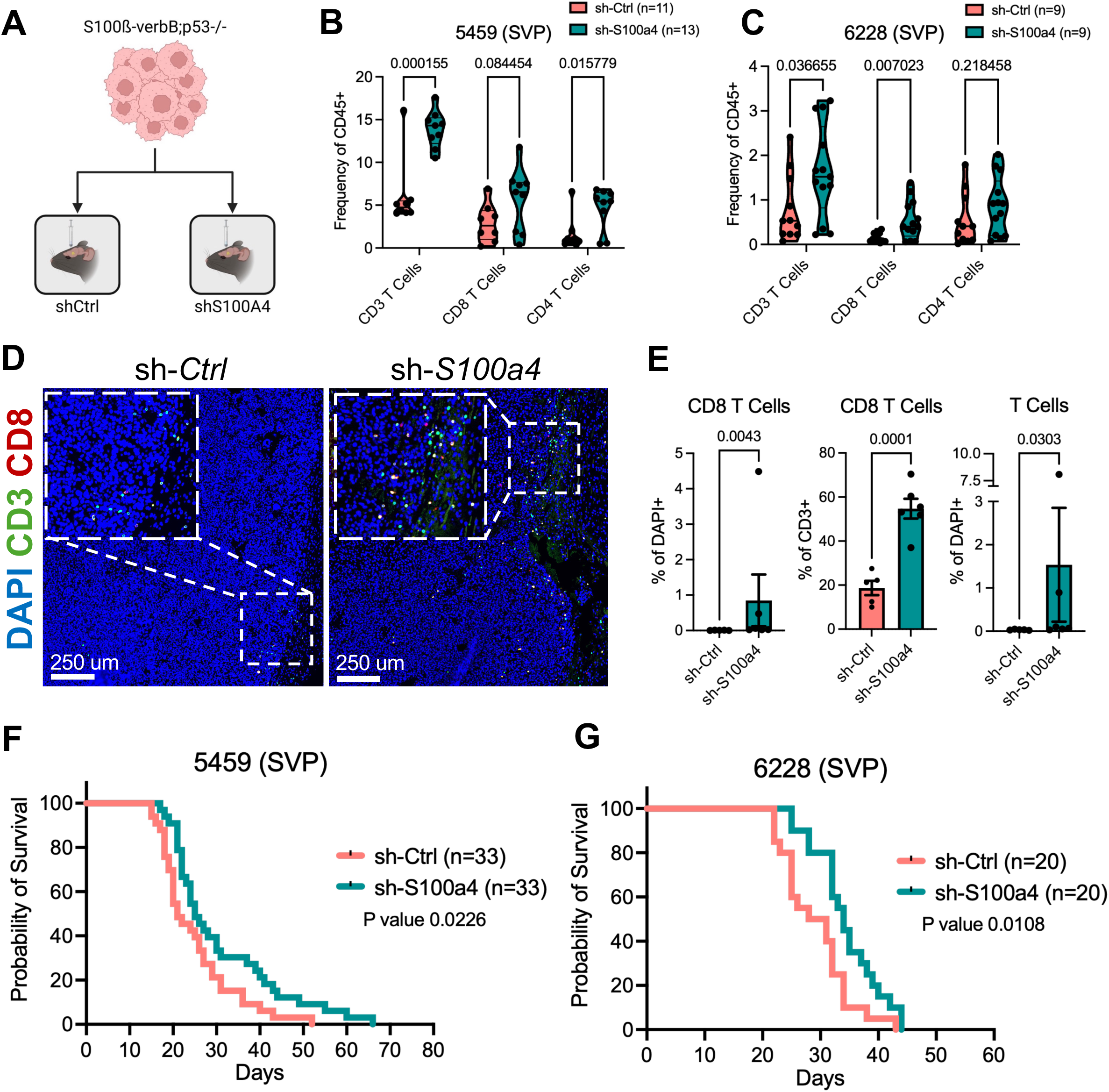
Knocking down *S100a4* in mouse glioma reprograms the GBM TME to promote T-cell infiltration. **A.** *In vivo* experimental schema for intracranial injection of control and S100a4-KD SVP model gliomaspheres into syngeneic B6 hosts. **B, C.** Flow cytometry analysis of intratumor immune cells from intracranially injected sh-*Ctrl* and sh-*S100a4* (B) 5459 gliomas or (C) 6228 gliomas. *p*-values determined by unpaired two tailed t-tests. **D.** Representative CD3 and CD8 immunofluorescence images of sh-*Ctrl* (n=5) or sh-*S100a4* (n=6) 5459 gliomas. **E.** Quantification of CD3 and CD8 T cells in sh-*Ctrl* and sh-*S100a4* 5459 gliomas. *p*-values determined by Mann Whitney test. **F, G** Kaplan-Meier survival curve of B6 mice intracranially injected with (F) 5459 gliomaspheres (n=33/group) or (G) 6228 gliomaspheres (n=20/group) with stable expression of sh-*Ctrl* or sh-*S100a4* shRNAs. *p*-values determined by logrank Mantel-cox test.

### *S100a4* knockdown sensitizes GBM to immunotherapy

While the survival benefit from *S100a4*-KD in glioma cells was consistent and significant, the effect size was modest. To understand the TME differences at end stage gliomas, we performed bulk RNA sequencing (RNA-seq) from control and *S100a4*-kD gliomas. Differentially expressed gene (DEG) analysis identified nearly 2600 genes with FDR <0.05 and LFC> 1 (**Figure 3A**). We analyzed the DEGs by conducting over-representation analysis (ORA), gene set variation analysis (GSVA), and network analysis, which consistently reported upregulation of T cell activation and pro-inflammatory pathways in *S100a4*-KD gliomas (**Figures 3B,C, Supplementary Figure 4A**). *S100a4*-KD significantly upregulated DEGs enriched in activated immune response-associated GO_BP gene sets, and significantly downregulated DEGs enriched in neural processes-associated gene sets (**Figure 3B**). We further showed through GSVA that *S100a4*-KD significantly upregulated genes associated with pro-inflammatory signaling pathways including IFNα and IFNy, as well as immune cell pathways including MHC, dendritic cells, and cytotoxic T cell pathways (**Figure 3C**). Immune cell deconvolution and abundance estimation analysis using the xCell package also showed increased immune cell populations, including CD8 T cells, CD4 T cells, B cells, NK cells, cDCs, and monocytes, supporting our IF and flow cytometry analysis (**Supplementary Figure 4B**).

**Figure 3:**
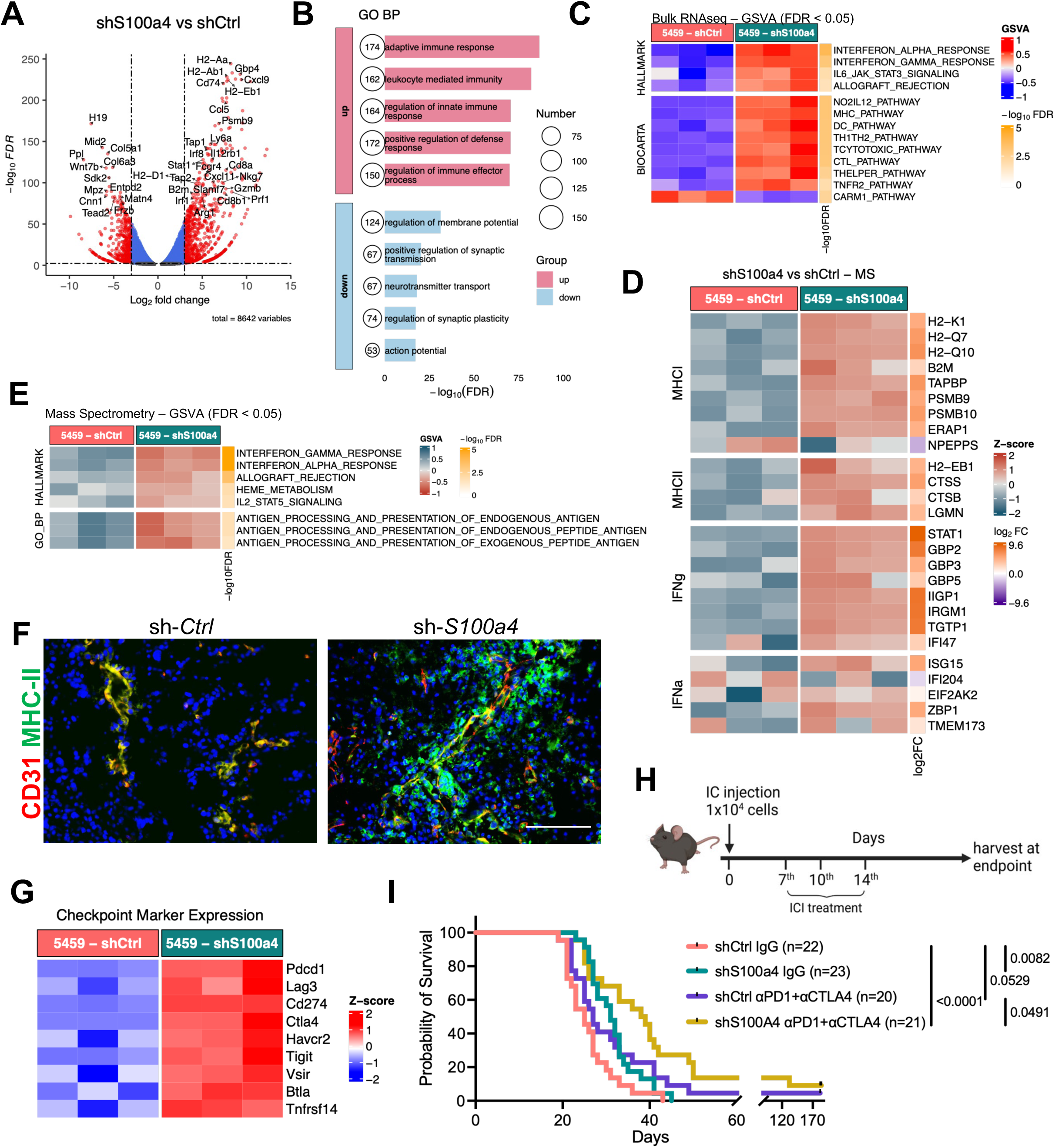
S100a4-KD activates Ifng signaling and antigen presentation to sensitizes mouse GBMs to immune checkpoint inhibitors. **A.** Volcano plot showing up and downregulated genes between sh-*S100a4* and sh-*Ctrl* 5459 tumors (n=3 per group). **B.** Over-enrichment analysis showing up and down regulated GO_BP gene sets for sh-*S100a4* tumors vs controls. **C.** GSVA of differentially enriched Hallmark and Biocarta gene sets for sh-*Ctrl* and sh-*S100a4* 5459 tumors. Adjusted *p*-value < 0.05 by False Discovery Rate correction. **D.** Heatmap showing normalized expression of select top differentially expressed proteins between sh-*Ctrl* and sh-*S100a4* 5459 tumors associated with MHC-I, MHC-II, Interferon-gamma, and Interferon-alpha pathway gene sets. **E.** GSVA of differentially enriched Hallmark and GO_BP gene sets for sh-*Ctrl* and sh-*S100a4* tumors from differentially expressed proteins identified via mass spectrometry analysis of tumor lysates (n=3 per group). Adjusted p-value < 0.05 by False Discovery Rate correction. **F.** Representative IF images of CD31 and MHC-II from sh-*Ctrl* and sh-*S100a4* 5459 tumors at endpoint. Scale bar represents 20 µm. **G.** Heatmap showing normalized expression of immune checkpoint molecules in shS100a4 5459 tumors from bulk RNAseq data. **H-I. (H)** Experimental schematic and **(I)** survival curve of ICI treatment in 5459 tumor bearing mice. *p*-values determined by log-rank Mantel-Cox test. n=20-23/group.

To measure changes at the protein level, we performed mass spectrometry analysis of control and S1004-KD gliomas. Consistent with the RNA-level analysis, the top enriched pathways were antigen processing and presentation (MHCI and MHCII)and interferon gamma and interferon alpha pathway proteins. (**Figures 3D, E, Supplementary Figure 4C**). Furthermore, immunofluorescence analysis of sh-*S100a4* vs sh-*Ctrl* tumors at endpoint further showed dramatically upregulated MHC-II expression, especially near tumor associated vasculatures (**Figure 3F**).

Despite increased anti-tumor immunity, *S100a4*-KD glioma-bearing mice died eventually (**Figure 2F, G**). This was not surprising given that S100A4 is also expressed by myeloid cells and T cells in the TME (*7*). To elucidate whether stromal S100A4 inactivates tumor infiltrating T cells in the *S100a4*-KD gliomas, we re-examined the bulk RNA-seq data and discovered that immune checkpoint molecules including *Pdcd1*, *Ctla4*, *Cd274*, *Tigit*, and *Lag3* (**Figure 3G**) were upregulated in *S100a4*-KD compared to controls. These results suggest that while *S100a4*-KD in glioma cells is sufficient to promote T cell infiltration, eventually the T cells become exhausted. On other hand, this observation suggests that S100A4 inhibition offers an opportunity to sensitize gliomas to ICI treatment.

To test this hypothesis, we injected sh-*Ctrl* or sh-*S100a4* 5459 gliomaspheres and treated them with either IgG controls or αPD-1+αCTLA-4, and measured survival (**Figure 3H)**. While αPD-1+αCTLA-4 only marginally improved survival compared to IgG treatment in control*-KD gliomas* (median OS: sh-*Ctrl*+ICI= 27 days vs sh-*Ctrl*+IgG=25 days, 8% increase in median OS, *p*-value=0.0529), *S100a4*-KD plus αPD-1+αCTLA-4 combination treatment significantly improved survival (median OS: sh-*S100a4*+ICI= 38.5 days vs sh-*Ctrl*+IgG= 25 days, 54% increase in median OS compared to control, *p*-value <0.0001) and resulted in several long-term survivors (**Figure 3I**). These results strongly suggest that S100A4 inhibition in glioma cells is sufficient to reprogram the TME and sensitizes GBM to ICIs.

### A novel αS100A4/αTFR bispecific antibody crosses the BBB and accumulates in GBM

To translate our findings for potential clinical application, we sought to develop an S100A4-blocking antibody that can cross the BBB. We previously reported an S100A4-blocking antibody that suppresses pre-metastatic niche formation in multiple *in vivo* mouse breast cancer lung metastasis models(*57*). However, treatment of 5459 glioma-bearing mice with the 4-11 monoclonal antibody (mAb) had no effect on survival**(Supplementary Figure 5A)**. Antibody therapeutics have difficulty crossing the blood-brain-barrier (BBB), and the 4-11 mAb may thus have limited ability to modulate S100A4 signaling in the GBM TME (*13, 14*). To overcome this physically relevant challenge, we designed a novel αS100A4/αTfR bispecific antibody (BsA), which fuses the bivalent S100A4 blocking antibody to a monovalent αTfR Fc fragment (**Figure 4A**). This design takes advantage of TfR-mediated transcytosis to bypass the BBB and enter the brain parenchyma (**Supplementary Figure 5B**)(*52, 53*).

**Figure 4:**
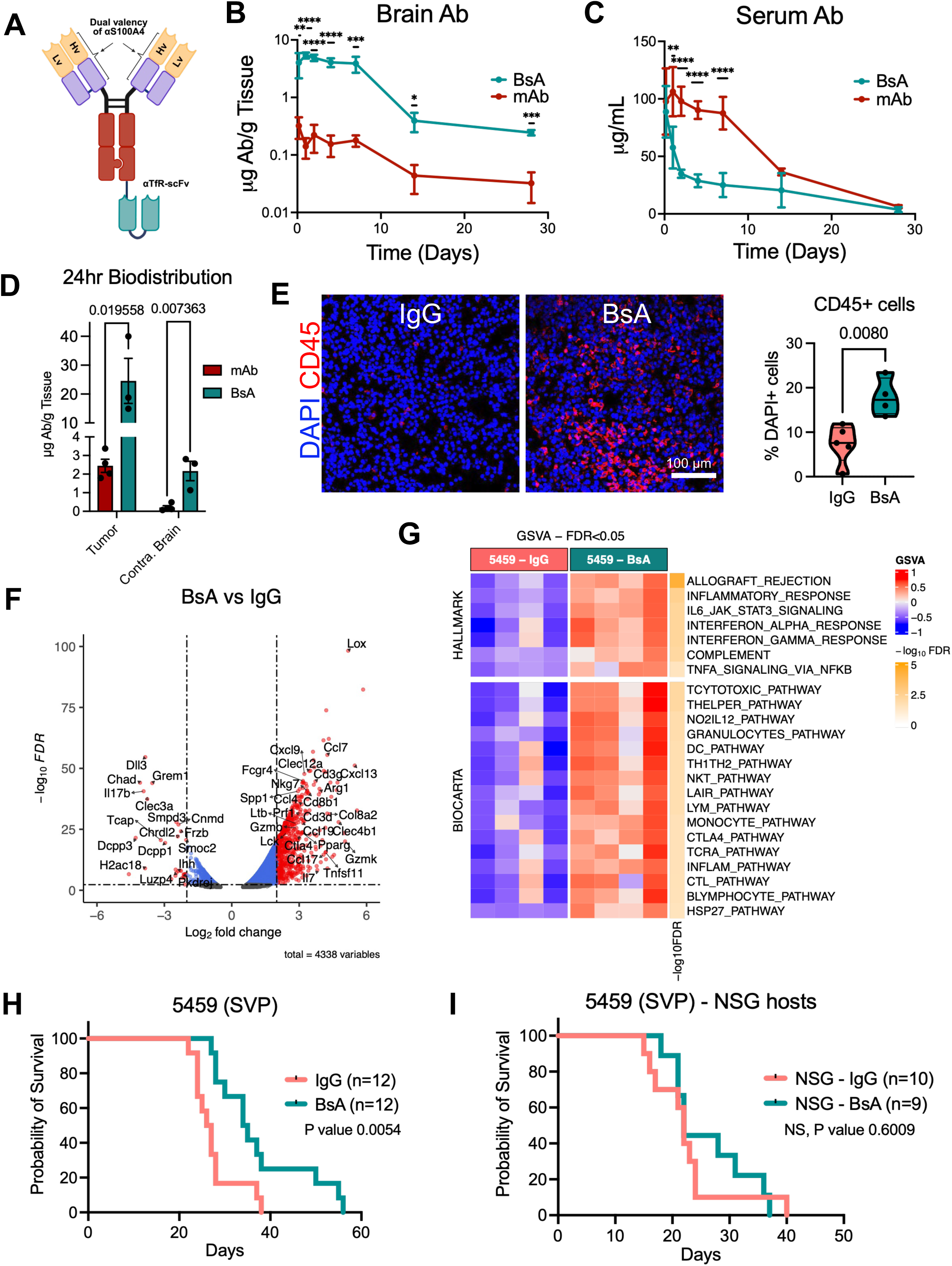
A bispecific S100A4/TFR antibody effectively penetrates the BBB and the brain tumor TME to reprogram the GBM TME and enhances survival in an immune dependent manner. **A.** Bi-valent S100A4 antibody. Hv: heavy chain variable region and Lv: light chain variable region of the anti-S100A4 antibody. **B,C**. BsA and mAb concentration in serum (B) and forebrain (C) post intraperitoneal injection (20mg/kg) as measured by ELISA. *p*-values determined by multiple unpaired two tailed t-test. (n=2-5/timepoint/group) *p<0.05, **p<0.01, ***p<0.001, ****p<0.0001. **D.** mAb and BsA concentration in tumor vs contralateral normal brain 24 hours post treatment as measured by ELISA. *p*-values determined by unpaired two tailed t-test. (n=3-4/group). **E.** Representative confocal IF images with Z stack of CD45+ cells in BsA and IgG treated 5459 tumors at endpoint and quantification of CD45+ cells in x5459 tumors at endpoint (n=4-5 per group). *p*-value determined by an unpaired two-tailed t-test. **F.** Volcano plot showing up and downregulated genes between IgG and BsA treated 5459 tumors at endpoint (n=4 per group). **G.** Gene set variation analysis (GSVA) of differentially enriched Hallmark and Biocarta gene sets between IgG and BsA treated 5459 tumors at endpoint. Adjusted p-value < 0.05 by False Discovery Rate correction. **H.** Kaplan-Meier survival curve of B6 mice intracranially injected with 5459 gliomaspheres and treated with BsA or IgG control. *p*-value determined by logrank Mantel-Cox test. (n=12/group). **I.** Kaplan-Meier survival curve of NSG mice intracranially injected with 5459 gliomaspheres and treated with BsA or IgG control. *p*-value determined by logrank Mantel-Cox test. (n=9-10/group).

We first assessed αS100A4/αTfR BsA biodistribution. We intraperitoneally (IP) injected non-glioma-bearing B6 hosts with αS100A4/αTfR BsA or αS100A4 mAb at a 20mg/kg dose and harvested brain and serum from the mice at various timepoints post-injection, starting from 4 hours up to 28D. We measured the αS100A4/αTfR BsA and αS100A4 mAb concentrations in brain lysates and serum using a sandwich ELISA (**Figure 4B, C**) and observed that BsA quickly accumulates in the brain parenchyma as early as 4 hours post-injection, at >10 times the concentration of the monoclonal antibody (**Figure 4C**). The BsA is maintained in the brain parenchyma at high levels for at least 7 days post-treatment. Brain BsA levels dropped significantly by D14 and D28, indicating a long half-life at around 8-9 days (**Figure 4C**). BsA serum levels peaked at around 4 hours after injection (mean 88.73 µg/mL), comparable to the mAb (mean 97.72 µg/mL), before dropping past 50% by 48 hours post-injection and gradually leveling down to between D3 and D14 post-injection, reaching similar levels to the mAb at D28 post injection (BsA 3.68 µg/mL vs mAb 6.29 µg/mL) (**Figure 4B**).

We next assessed the BsA and mAb tumor uptake in 5459 glioma-bearing mice, 24 hours post-treatment (20 mg/kg, IP). We stained for the BsA in SVP glioma and contralateral normal brain tissues via IF, using secondary antibodies against Human IgG. There were easily detectable BsA levels but not the mAb in glioma tissues (**Supplementary Figure 5C**). To quantitate these differences, we generated lysates from tumor and contralateral brain from 5459 glioma-bearing mice (n=3-4) and measured the BsA and mAb concentrations using a sandwich ELISA. Consistent with our IF findings, the BsA accumulates in the glioma at approximately 10x the monoclonal (BsA 24.56 µg/g tumor tissue vs mAb 2.44 µg/g tumor tissue, *p*-value=0.019558) (**Figure 4D**). As antibody therapeutics may lead to a host immune response, including anti-drug antibodies production (ADA), we conducted flow cytometry of splenocytes and circulating PBMCs from D28 B6 mice after one-time BsA treatment, and found no differences compared to mAb treated control populations (**Supplementary Figure 5D**). Similar to the mAb(*57*), the BsA was well tolerated and no changes in weight loss was detected.

### αS100A4/αTfR BsA remodels the GBM TME and sensitizes mouse GBMs to immune checkpoint inhibitors

As *S100a4*-KD in glioma cells or *S100a4*-deletion in the host remodels the GBM TME to a more pro-inflammatory state(*7*) (**Fig 3B, C**), we hypothesized that BsA treatment will have a similar immune modulatory effect. We injected 5459 SVP glioma cells into B6 hosts and treated them from day 7 post-injection, once weekly, based on our PK study (**Figure 4C**). We first analyzed the overall immune landscape via IF and detected increased CD45+ cells in BsA treated gliomas vs control IgG treated (**Figure 4F)**. We then analyzed BsA vs IgG treated gliomas via bulk RNA sequencing (**Figures 4G**). Similar to *S100a4-*KD gliomas, DE, ORA, and GSVA analyses indicate significant enrichment of pro-inflammatory signaling pathways such as TCYTOTOXIC PATHWAY, THELPER PATHWAY, TCRA PATHWAY, INFLAMMATORY RESPONSE, INTERFERON GAMMA RESPONSE, INTERFERON ALPHA RESPONSE, and TNFA SIGNALING VIA NFKB (**Figures 4G, Supplementary Figure 6A**), as well as increased expression of genes associated with cytotoxic, helper T cell, and overall adaptive immune response pathways, such as *Cd3d*, *Cd8b1/a2*, *Gzmb*, *Gzmk*, *Il7*, *Tnfsf11/Rankl*, *Ccl7*, *Clec4ba*, *Clec12a*, *Cxcl9*, *Nkg7*, and others (**Figure 4G, Supplementary Figure 6B**) in BsA treated gliomas. Importantly, BsA treatment significantly improved survival of glioma-bearing hosts compared to IgG controls (IgG control: median OS=26.5 days; BsA median OS=34.5 days, 30.2% increase in median OS, *p*-value=0.0054) (**Figure 4H**). To determine whether the survival benefit is provided by immune reprogramming, we injected gliomas cells into immune-deficient NOD.Cg-*Prkdc^scid^ Il2rg^tm1Wjl^*/SzJ (NSG) mice, which lack functional B, T, and NK cells and dysfunctional myeloid cells (**Figure 4I**). BsA treatment survival benefit observed in B6 hosts was abolished in NSG hosts, indicating that survival benefit was dependent on the intact immune system (**Figure 4I)**. Overall, our results indicate that αS100A4/αTfR BsA treatment can provide similar immune modulatory effects and survival benefit as genetic *S100a4*-KD.

### BsA sensitizes GBM to ICI treatment and induces TLS-like features

Genetic knockdown of *S100a4* reprograms the GBM TME to a pro-inflammatory state (**Figures 3A-E**) but T cells become exhausted over time (**Figure 3G**); however, this sensitizes them ICI treatment(**Figure 3I**). To determine whether the BsA treatment similarly reprograms the TME, we examined immune checkpoint molecule expression in BsA treated gliomas, and detected similar upregulation of immune checkpoints, including *Pdcd1, Cd274,* and *Ctla4* (**Figure 5A**). Therefore, we hypothesized that BsA plus ICI combination therapy may be a clinically translatable approach. To test this hypothesis, we combined weekly BsA 20 mg/kg treatment with bi-weekly αPD-1 10mg/kg treatment (Combo) in 5459 tumor-bearing B6 mice starting at 7 days post tumor injection (**Figure 5B)**. The Combo treatment nearly doubled the median survival compared to IgG control (median OS for Combo 48 days vs IgG 26.5 days, 81.1% increase, *p*-value<0.0001) and significantly longer than either αPD-1 or BsA monotherapy alone (**Figure 5C)**. Analysis of intratumor immune cell infiltrates via flow cytometry showed that the Combo treatment led to a significant increase in tumor reactive CD8 T cells (CD39+CD8+PD1+) (**Figure 5D, Supplementary Figure 6C**). We further assessed intratumor immune cell infiltrates via IF and observed a significant increase in T cell infiltration in Combo treated gliomas (**Figures 5E, F, Supplementary Figures 7A, B**). The BsA+αPD-1 Combo treatment significantly increased CD3+ T cells, CD4+ T Cells and CD8+ T cells, including Perforin-1(PRF1)+ CD8 T cells, which appeared primarily at the tumor border (**Figure 5E, F, Supplementary Figures 7A, B**). To further assess the spatially organized phenotype of T cell infiltration in BsA+αPD-1 treated tumors, we conducted multiplex immunofluorescence (COMET) on treated samples (**Figure 5G)**. We first compared immune infiltrates in tumor core vs tumor border, including CD4 and CD8 T cells, for each treatment arm. There were no notable differences in IgG and αPD1 treated gliomas. However, in BsA treated and the combination treated gliomas, immune infiltrates increased in the tumor border regions, and T cell infiltration was notably increased in the tumor border regions for the combination treated (**Figure 5G, H, Supplementary Figure 7C**). Quantitation of immune cells, T cells, and macrophages at tumor core vs periphery show dramatic increase in CD45+, CD8+PD1+, CD8+TBET, and CD4+FOXP3+ cells at the tumor border in BsA or BsA+αPD1 Combo treated tumors (**Figures 5G,H, Supplementary Figure 7C**). In addition, we observed increased proliferation of T and B cells in the cervical lymph node of BsA+αPD-1 Combo treated tumors at 2 weeks post-treatment, suggesting an increased anti-tumor immune response (**Supplementary Figure 8**).

**Figure 5:**
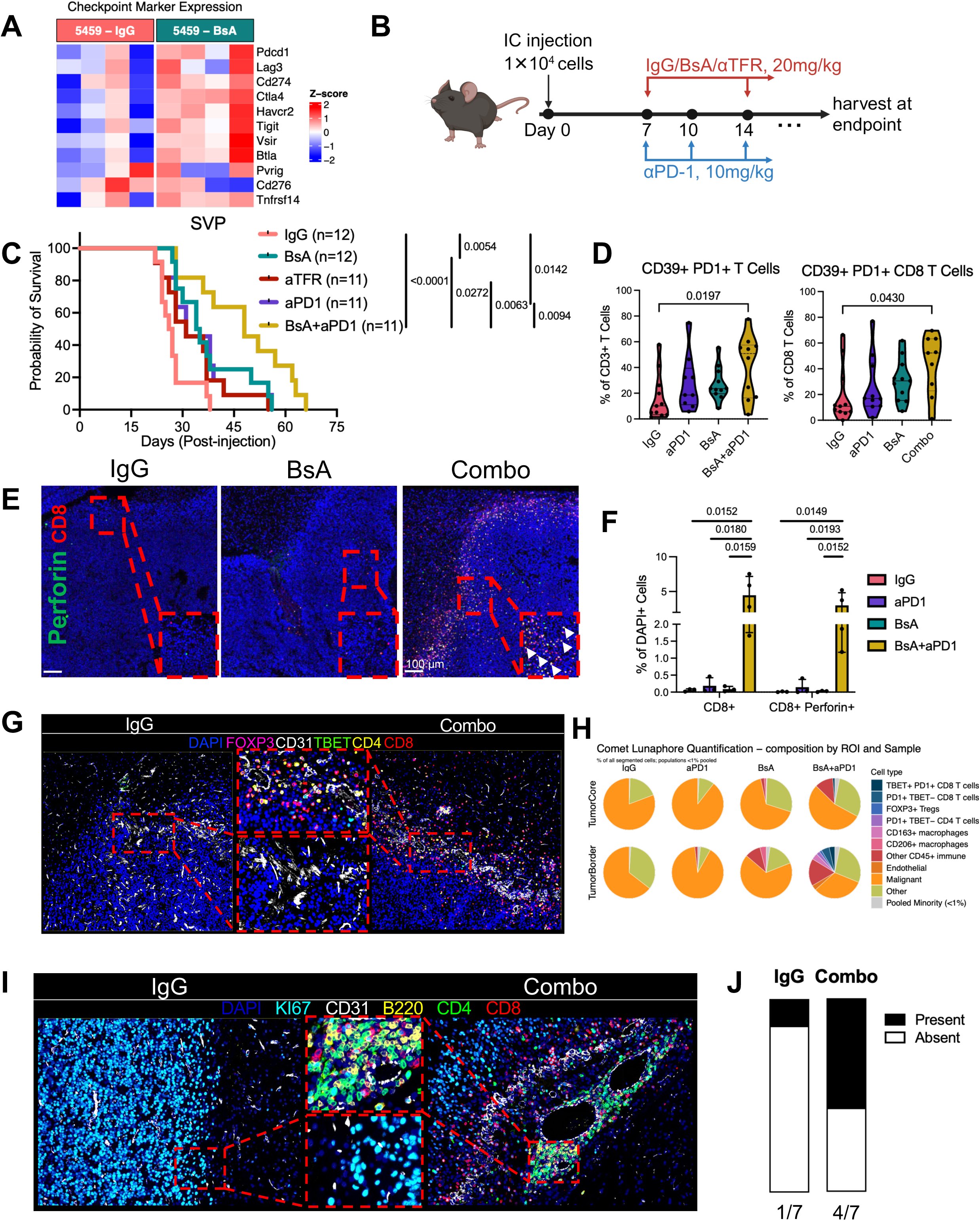
BsA treatment synergizes with immune checkpoint inhibitors to significantly enhance survival. **A.** Heatmap showing normalized differential expression of immune checkpoint molecules in IgG and BsA treated 5459 tumors from bulk RNAseq data. **B.** Experimental schema for *in* vivo survival study. B6 mice were intracranially injected with 10,000 dissociated SVP (5459) gliomaspheres. Starting at day 7, mice were treated with IgG, BsA, or αTFR 20mg/kg weekly or αPD-1 10mg/kg twice weekly. BsA+αPD-1 treatment group received weekly BsA 20mg/kg treatment and twice weekly αPD-1 10mg/kg treatment. All treatments continued until endpoint. **C.** Kaplan-Meier survival curve of B6 mice intracranially injected with SVP gliomaspheres and treated with IgG (median OS: 26.5), BsA (median OS: 34.5), αTFR (median OS: 31), αPD-1 (median OS: 31), or BsA+αPD-1 (Combo) (median OS: 48). *p*-values determined by logrank Mantel-cox test. n=11-12/group. Data from Figure 4E is reproduced here (same experiment) is reproduced here. **D.** Spectral flow cytometry analysis with violin plots showing percentage of CD39+ PD1+ CD3 T cells and CD8 T cells from endpoint tumors (n=9-11 per group). Cells are gated off LiveDeadBlue-CD45+ CD11b-CD3+ and LiveDeadBlue-CD45+ CD11b-CD3+ CD8+ populations. *p*-values determined by one way ANOVA with Dunnet correction for multiple comparisons, p < 0.05 displayed. **E.** Representative IF images for CD8 and Perforin from IgG, BsA and Combo treated endpoint SVP samples. Scale bars represent 100 µm. Arrows indicate CD8+ Perforin+ cells. **F**. Quantification of IF analysis from endpoint SVP tumors. n=3-4 per group. *p*-values determined by one-way ANOVA with Šidák correction for multiple comparisons, p < 0.05 displayed. Bar plots and error bar indicate mean and SEM, respectively. **G.** Representative COMET assay IF plots for IgG and Combo treated tumors. **H.** Pie chart showing proportions of immune cell populations in tumor core and tumor border regions of treated SVP tumors from COMET assay, grouped by treatment group. N=2/group. **I.** Representative COMET assay IF plots for IgG and Combo treated tumors showing presence of tertiary lymphoid structures in Combo treatment samples. Arrows indicate TBET+ CD8+ cells. **J.** Parts-of-whole vertical slice plot showing proportion of samples with TLS-like structure present, n=7/group.

Intriguingly, we discovered aggregates of CD4 T cells, CD8 T cells, and B cells in Combo treated gliomas (**Figure 5I**) that resembled early tertiary lymphoid structures (TLS). These TLS-like structures lacked classical germinal centers but were B and T cell rich, suggesting active adaptive immune response. The presence of TLS has been observed in both mouse and human gliomas and is positively correlated with both prognosis and immunotherapy response (*30, 34–36*). We quantified TLS-like structures by defining it as cellular aggregate of at least 10 B cells and 20 T cells clustered within a 900,000 µm^2^ region. We thus stained for CD8 and CD19 in IgG and Combo treated samples and observed TLS-like structures in 4 of 7 samples analyzed vs 1 of 7 samples in IgG control gliomas (**Figure 5J**), indicating preferential induction of TLS-like structures with BsA+αPD1 Combo treatment.

### Spatial transcriptomics analysis support TLS formation in Combo treated gliomas

To take an unbiased approach, we conducted Visium spatial transcriptomics analysis of Combo and control treated gliomas (**Figure 6A**). We first identified tumor regions vs normal brain regions using H&E images and inferred copy number alteration (CNA) analysis, which we validated with cell type deconvolution analysis (**Supplementary Figure 9**). We then conducted nNMF to identify consensus gene metaprograms expressed across tissues regions from all treatment arms and annotated them based on ORA of top expressed metaprogram genes (**Figure 6B, C Supplementary Figure 10A-C**). For example, we identified and annotated metaprograms associated with cell proliferation (Proliferative), antigen presentation and interferon gamma signaling (AP-IFN), and hypoxia-induced signaling (Hypoxia), among others (**Figures 6B-D, C, Supplementary Figures 10A-D**). We calculated metaprogram module scores, assigned each spot to its dominant metaprogram, and classified metaprogram modules as malignant or non-malignant based on the spot level malignancy composition (**Figures 6B, C**). Metaprogram assignment analysis by treatment arm and tissue region determined by dominant MP score per spot (**Figures 6B, C**) or by mean MP signature score per spot (**Figure 6D**) showed striking differences in the tumor core and border. In the tumor core, Combo treated gliomas stood out as having highly enriched AP-IFN module score, whereas all other treatment group spots showed similar module scores, including positive module scores for Proliferative/ECM and Hypoxia metaprograms (**Figures 6C, D**). Similarly, the tumor border regions in BsA and Combo treatment were more dominated by the AP-IFN metaprogram, whereas αPD1 had the highest enrichment of the inflammatory module, and IgG and αPD-1 had a much more diverse metaprogram assignment including non-malignant metaprograms (**Figures 6C, D**). We validated this by assessing expression of antigen presentation associated genes at the spot level, split by tumor region and normal brain, and found that both MHC-I and MHC-II associated genes were dominantly expressed in both the tumor border and tumor core of Combo treatment gliomas compared to other treatment arms (**Supplementary Figure 10E**).

**Figure 6:**
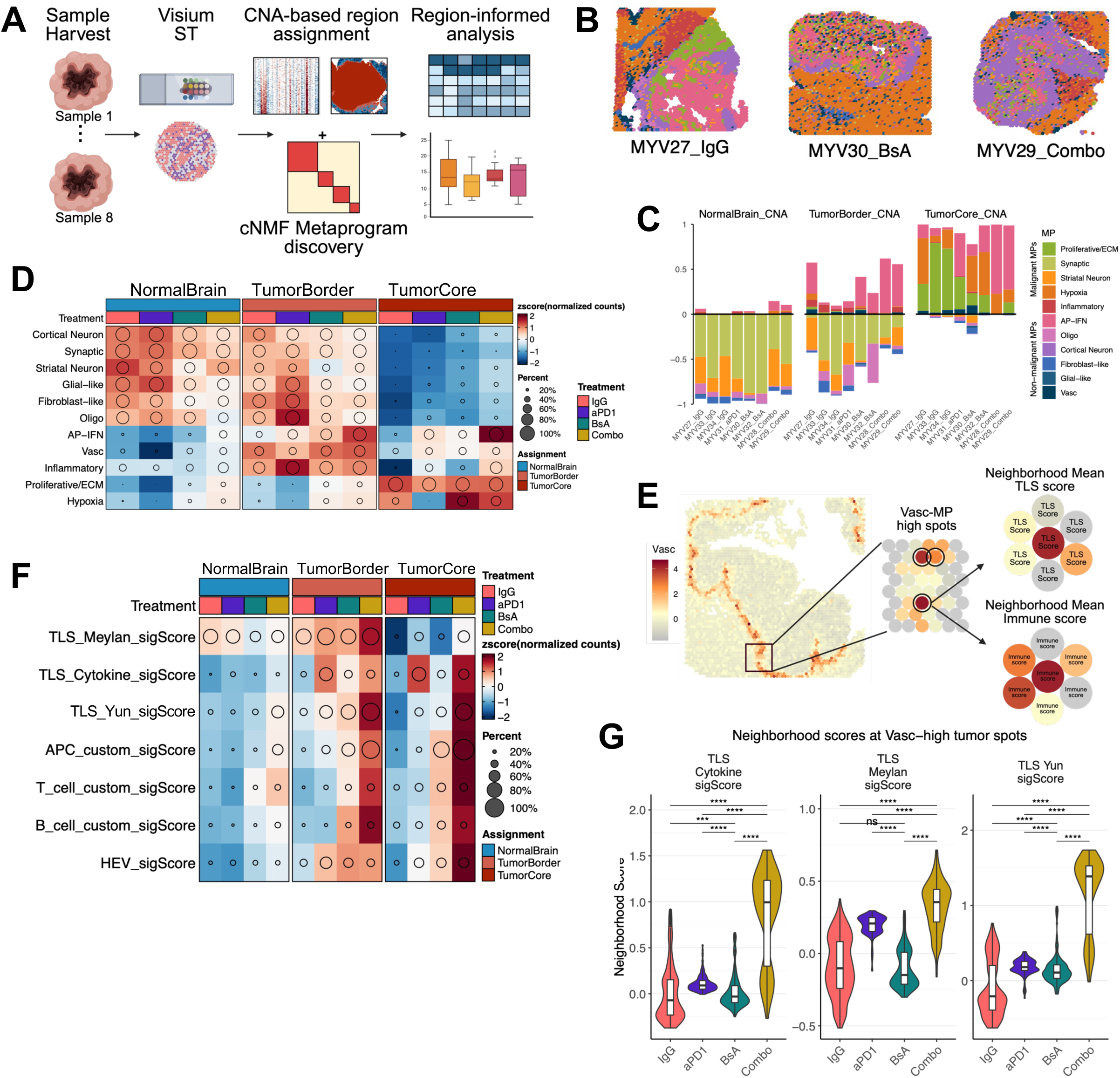
BsA synergizes with aPD-1 treatment to induce antigen presenting hubs at the vascularized tumor border that form TLS-like structures. **A.** Graphic for Visium spatial transcriptomics analysis workflow. **B.** Spatial dimension plot showing spot level dominant nNMF MP. Each spot is assigned a dominant nNMF MP based on top signature score. Spots are colored by assigned nNMF MP. **C.** Stacked bar plots showing spot level proportion of metaprograms for each sample, split by assigned tissue region. **D.** GroupHeatmap showing mean row-z-scored nNMF MP module scores grouped by treatment group and split by tissue region assignment. **E.** Schematic workflow showing neighborhood score calculation of TLS and immune cell scores in spots expressing high Vasc nNMF MP signature scores (Vasc-high) in the tumor core and tumor border. **F.** GroupHeatmap showing mean row-z-scored TLS and cell type module scores grouped by treatment group and split by tissue region assignment. **G.** Violin plot showing neighborhood scores of published and custom TLS gene modules in Vasc-high expressing spots from the tumor border and tumor core, grouped by treatment group. Box represents median and interquartile range. Vasc-high spots were determined taking all tumor border and tumor core spots from all samples, computing a signature score from Vasc nNMF MP genes, and thresholding at 2 standard deviations above the average.

Our COMET assay analysis also indicated increased endothelial cell abundance at the tumor border compared to the tumor core in Combo treated samples (**Figure 5F Supplementary Figure 7C)**. To explore this further, we stained all treatment group gliomas with anti-CD31 antibody and quantified vasculature areas in the tumor core and tumor border (**Supplementary Figures 11A, B**). While there was no difference in the vascular density in tumor cores, vasculature density was significantly higher at the tumor border in BsA and Combo treated gliomas (**Supplementary Figures 11A, B**). This phenotype was also observed in *S100a4*-KD gliomas (**Supplementary Figures 11C, D**), suggesting that S100A4 inhibition results in increased angiogenesis and T cell infiltration at the tumor border.

Since increased T cell infiltration was proximal to the vasculature at the tumor border in Combo treatment, we hypothesized that the vasculature formed at the tumor border may be high endothelial venule (HEV)-like vessels that improve T cell trafficking into those regions. To answer this question, we turned to our spatial transcriptomic data. We first scored spots for gene modules for T cells, B cells, and HEVs using a panel of curated signature genes (**Supplementary Table 1, Figure 6E**). To identify spots enriched for tumor vasculature, we subset spots in Tumor Border and Tumor Core that scored two standard deviations above the mean for Vasc MP module scores (**Figure 6E**). For each of these Vasc-high spots, we then calculated a neighborhood score for B cells and T cells. To define the neighborhoods, we took each Vasc-high spot and its neighboring spot within a one-spot radius and calculated the mean spot module score for HEVs and immune cells (**Figure 6E**). We then directly compared these neighborhood scores by treatment arm and conducted correlation analysis between neighborhood HEV and T cell scores (**Supplementary Figure 11E, F**). Neighborhood B and T cell scores were higher in Vasc high spots from Combo treatment samples compared to controls (**Supplementary Figure 11E**). HEV scores and neighborhood T cell scores in Vasc-high spots were positively correlated with each other, while both scores were preferentially higher in Vasc-high spots from Combo treatment samples compared to controls (**Supplementary Figure 11F**). Our results thus indicate that Combo treatment samples induced HEVs that correlated with increased B and T cell infiltration.

Using a similar rationale, we scored our Visium samples for TLS scores derived using curated and published gene sets(*58, 59*) (**Supplementary Table 1**). We found that spot level TLS scores and neighborhood TLS scores in Vasc-high spots were also higher in Combo treatment compared to controls (**Figure 6F, G**). These findings support the notion that αS100A4/αTfR BsA treatment in combination with ICIs can induce TLS formation through HEVs and increased T cell infiltration and activation.

### S100A4 inhibition benefits all molecular subtypes of GBM

Given that human glioma is highly heterogenous, with different molecular subtypes, genetic drivers, and TME compositions that affect treatment outcomes, we asked whether inhibition of S100A4 can extend survival and reprogram the GBM TME across different GBM molecular subtypes. To answer this question, we took advantage of a new cohort of mouse glioma models representing different molecular subtypes of GBM: EIP: *tetO-EGFRviii; Ink4a/Arf^-/-^ ; Pten^-/-^* model (classical)(*60*), fPPP: *mGFAPcre; p53^flox/flox^; Pten^flox/flox^; PIK3CAH1047R^flox/+^* (mesenchymal), KAB: *N/tv-a; PDGFA; shp53* (pro-neural)(*61*), and KABN: *N/tv-a; PDGFA; shp53; shNF1* (pro-neural/mesenchymal). We isolated primary gliomaspheres from spontaneous gliomas from each of these models and confirmed that they represent human GBM heterogeneity in glioma cells and the TME (Tran et al., in preparation). We knocked down *S100a4* in each line, and intracranially injected sh-*Ctrl* and sh-*S100a4* gliomaspheres into syngeneic B6 hosts. *S100a4-KD* significantly extended host survival across all mouse models tested (**Figures 7A-D)** (EIP: median OS sh-*S100a4* 89 days vs sh-*Ctrl* 59 days, 50.8% increase in median OS, *p*-value=0.0051; fPPP: median OS sh-*S100a4* 25 days vs sh-*Ctrl* 21 days, 19% increase in median OS, *p*-value=0.0047; KAB: median OS sh-*S100a4* 53.5 days vs sh-*Ctrl* 43 days, 24.4% increase in median OS, *p*-value=0.0165; KABN: median OS sh-*S100a4* 57 days vs sh-*Ctrl* 43 days, 32.6% increase in median OS, *p*-value=0.0275). We also assessed the immune landscape of sh-*Ctrl* and sh-*S100a4* gliomas via flow cytometry and IF. Consistent with the SVP model (**Figure 2**), *S100a4*-KD significantly increased intratumor immune infiltrate and activation, including CD8 T cells in other models (**Figure 7E-G**).

**Figure 7:**
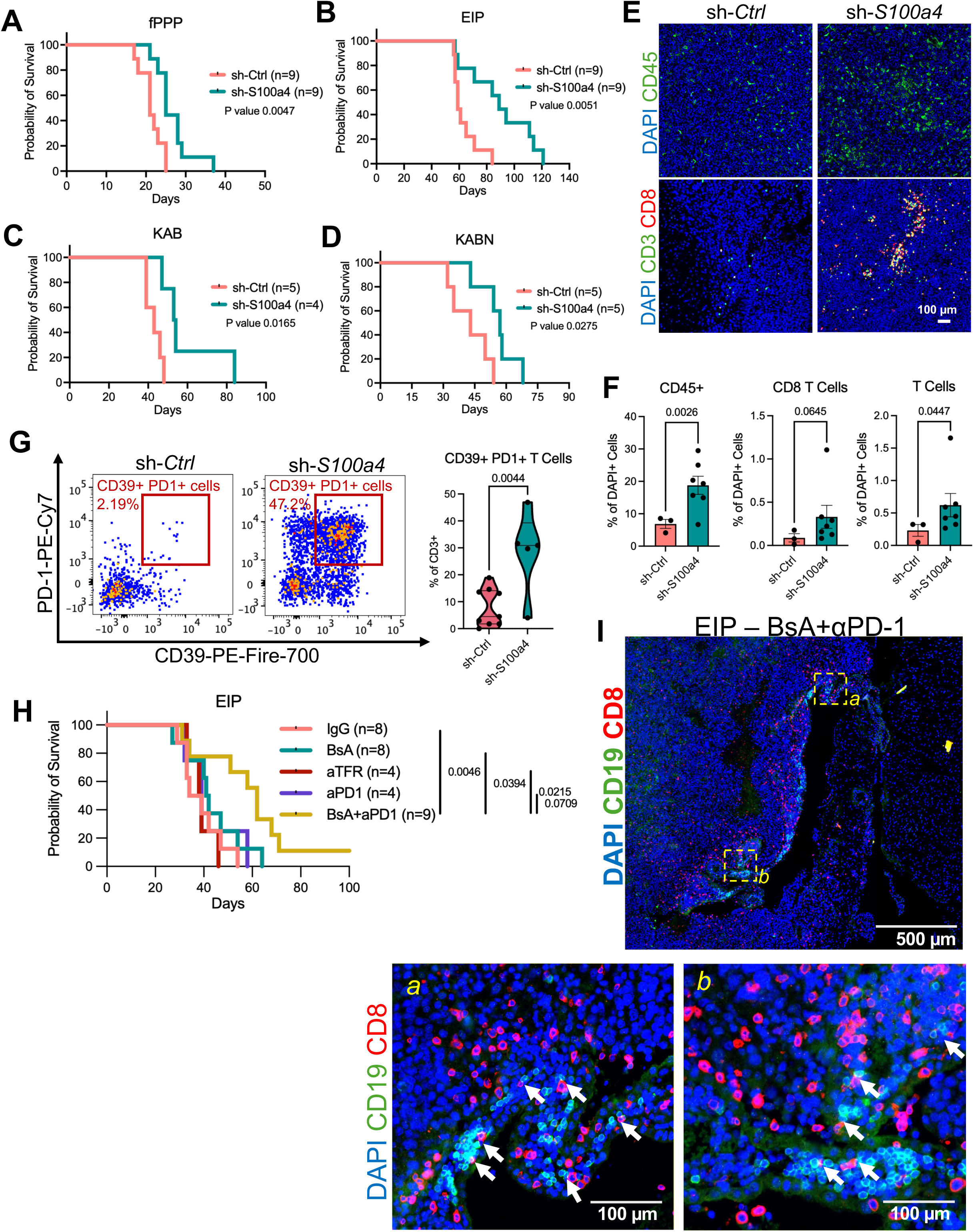
S100a4-KD extends survival and enhances T cell infiltration in multiple mouse GBM models that faithfully model the human subtypes of GBM. A-D. Kaplan-Meier survival curve of B6 mice intracranially injected with sh-*Ctrl* and sh-*S100a4* (A) 3422 (fPPP model) gliomaspheres, (B), 2225 (EIP model) gliomaspheres, (C) KAB203 (KAB model) gliomaspheres, and (D) KAB200 (KABN model) gliomaspheres. **E.** Representative IF images of CD45, CD3, and CD8 co-staining in sh-*Ctrl* and sh-*S100a4* 3422 (fPPP) tumors. Scale bar represents 100 µm. **F.** Quantification of CD45+, CD3+, and CD3+ CD8+ T cells in sh-*Ctrl* (n=3) and sh-*S100a4* (n=7) 3422 gliomas. *p*-values determined by Welch’s t-test. **G.** Flow cytometry analysis of intratumor T cells in sh-*Ctrl* (n=9) and sh-*S100a4* (n=5) 3422 gliomas at endpoint, with quantification. *p*-values determined by unpaired two-tailed t-test. Cells are gated off LiveDeadBlue-CD45+ CD11b-CD3+ populations (Ctrl: N=9, KD: N=5). **H.** Kaplan-Meier survival curve of B6 mice intracranially injected with 2225 (EIP model) gliomaspheres and treated with IgG (n=8), BsA (n=8), aTFR (n=4), αPD-1 (n=4), or BsA+ αPD-1 (Combo)(n=9). *p*-values determined by logrank Mantel-Cox test. **I.** Representative IF plot displaying aggregate of CD8+ and CD19+ showing a TLS-like structure with arrows indicating B-T cell interaction.

To assess the generalizability of the BsA treatment, we assessed the BsA+αPD-1 combination treatment efficacy in another glioma model (**Figure 7H**). The combination therapy significantly extended survival compared to IgG control (median OS for Combo 62 days vs IgG 36.5 days, 69.9% increase, *p*-value=0.0046), while we did not see a significant survival benefit with BsA or αPD-1 alone vs IgG control (**Figure 7H**). Consistent with the SVP model (**Figure 5I**), we observed that a majority of Combo treated EIP samples assessed (3/4) had the presence of B/T cell aggregates resembling TLS-like structures (**Figure 7I**).

Overall, these findings support the concept that targeting S100A4 with or without ICIs can improve survival across heterogenous mouse models representing different subtypes of human glioma. Mechanistically, S100A4 inhibition combined with ICI inhibition induces TLS formation and increases T and B cell infiltration and activation, leading to improved survival of glioma-bearing mice. Thus, S100A4 represents a novel immunotherapy target for treating glioma, and the αS100A4/αTfR BsA warrants further investigation for clinical translation.

## DISCUSSION

Current immunotherapies including immune checkpoint blockade (ICB) and chimeric antigen receptor T (CAR-T) cell therapies have had limited efficacy in treating GBM patients (*2–4, 14, 62, 63*). Clinical trials with neoadjuvant anti-PD-1 treatment in recurrent GBM patients have had mixed results, including an early study that indicated clinical benefit, which was not reproduced in a subsequent study(*63, 64*). Recent studies assessing the subset of patients responsive to immunotherapies suggest several molecular features that may predict clinical benefit with these modalities(*63, 65–67*). For example, in a study assessing anti-PD-L1 treatment in newly diagnosed GBM patients, patients with IFN-γ signaling and immune enrichment in pre-treatment tumor samples had improved response to PD-L1 treatment(*65*). A more recent study suggests that mesenchymal GBM signatures are associated with improved ICB outcomes, including pre-treatment tumors with higher HLA class I expression and T cell infiltration(*66*). Overall, these studies indicate immunotherapy response in GBM depends on GBM TME prior to treatment, similar to other cancers. Successful ICB treatment requires a T cell-rich microenvironment with sufficient antigen presentation, either by myeloid cells or by tumor cells. Therefore, strategies that reverse the immunosuppressive TME and improve T cell infiltration may be pre-requisite for successful deployment of immunotherapies in GBM.

Here, we report that genetic or pharmacological targeting of S100A4 is sufficient to reprogram the GBM TME and sensitize tumors to ICB. Both glioma and immune cells express and secrete S100A4 to promote an immune suppressive TME. We previously published that genetic loss of S100A4 in stromal cells is sufficient to increase T cell activation and macrophage phagocytosis, resulting in a significant survival benefit (*7*). Here, we show that genetic knockdown of S100A4 in glioma cells alone is also sufficient to extend survival in 5 different glioma models, although the effect size is smaller than with TME deletion. To translate these findings, we developed an S100A4 blocking antibody (*57*). However, as anticipated, the monoclonal antibody showed limited penetrance through the BBB. To overcome this well-known barrier, we developed a novel bispecific antibody targeting S100A4 and TfR, which utilizes TfR-mediated transcytosis to bypass the BBB and enter the brain parenchyma and tumor tissues. The αS100A4/αTFR BsA accumulates at high levels in CNS tissues compared to the monoclonal antibody, and at 10x higher concentrations in tumor tissues compared to contralateral normal brain (**Figure 4A-D**).

Clinically, human GBMs can be stratified into the classical, pro-neural, and mesenchymal molecular subtypes based on their molecular profiles at the bulk tissue level(*68*). Molecular subtype classification influences both the TME immune composition as well as therapy response, including immunotherapy(*65, 69, 70*). Using any individual pre-clinical mouse GBM model fails to capture this vast diversity. Here, we tested *S100a4*-KD and BsA treatment efficacy in multiple mouse models with various genetic drivers that reflect the different human molecular subtypes. Survival benefit magnitude varied by mouse model, suggesting that potential clinical translation will have similarly heterogenous effects. Reflective of this varied survival benefit observed in mouse models, our analysis of the GLASS dataset indicates that amongst IDH-wildtype glioma patients, *S100A4* expression significantly stratifies survival in recurrent IDH-wildtype glioma patients with mesenchymal-subtype, but not other molecular subtypes. In concordance, we see robust survival benefit with *S100a4*-KD+ICI and BsA+ICI treatment in the SVP model, which models mesenchymal-subtype GBMs. Our results warrant further investigation of S100A4 as a clinical target in mesenchymal-subtype GBM and nominate the S100A4/TFR BsA as a potential therapeutic for this devastating disease with limited treatment options. Additionally, GLASS dataset analysis indicates that *S100A4* expression significantly stratifies survival in patients with IDH-mutant gliomas, and S100A4 expression increases in IDH-mutant gliomas at recurrence. Thus, S100A4 may also be a clinical target in IDH-mutant gliomas to prevent progression and/or recurrence.

Our analysis indicates that BsA+ICI treatment results in accumulation of high endothelial venules (HEVs) at the tumor border and the formation of TLS-like features with aggregates of T and B cells. While the presence of HEVs in human glioma remains controversial (*34, 71*), TLSs have been associated with better ICI response in multiple cancer types (*31–33*). TLS-like structures with varying maturation stages have been reported in murine and human GBM, and their presence correlates with OS (*34, 35*). Formation of different subtypes of TLS has been observed in glioma patients; the TLS-like structures formed in mouse gliomas (SVP and EIP models) with BsA+αPD-1 treatment are CD8 T-cell rich, and includes many B cells, which are rare in untreated GBMs. In previous reports in human glioma, TLS subtypes associated with high B and T cell interactions are characterized by expression of genes related to antigen presentation (*34*). In concordance, antigen presentation is significantly enriched in both *S100a4*-KD and BsA+αPD-1 treated samples as well as *S100a4-*KD gliomas. Overall, we propose a framework where genetically or therapeutically targeting S100A4 sensitizes GBMs to ICI to increase T cell infiltration and activation, leading to a more pro-inflammatory antigen presentation rich TME amenable to ICB.

S100A4 is also a therapeutic target in modulating fibrotic diseases, with multiple ongoing clinical trials evaluating the therapeutic efficacy of monoclonal S100A4 antibodies in psoriasis and pulmonary fibrosis. Favorable safety, tolerability, pharmacokinetic, and immunogenicity profile of CAL101/AX-202 S100A4 targeting antibody in a Phase 1 trial (NCT05965089) has led to the advancement of CAL101 to Phase 2 clinical trial (NCT06736990) for idiopathic pulmonary fibrosis(*72*). Multiple bispecific antibodies utilizing agonistic TFR antibodies for CNS delivery are currently in clinical trials for Alzheimer’s disease(*73, 74*). Combined with our promising pre-clinical data, these studies strongly support clinical development of the αS100A4/αTFR BsA to treat GBM.

This study provides robust pre-clinical evidence that targeting S100A4 reprograms the GBM TME and sensitizes GBM to immunotherapies. It will be interesting to test whether S100A4 inhibition can improve the efficacy of other immunotherapies, such as CAR-T that requires significant infiltration of T cells into the GBM TME. It is well documented that the BBB in GBM is partially disrupted, but overall remains an obstacle for GBM therapeutics including immune checkpoint inhibitors and adoptive cellular therapies like CAR-T(*75*). CAR-T and other therapies such as oncolytic viruses are increasingly administered directly locally(*21, 76*) but their spread or activation state within the highly immune suppressive TME may continue to pose a challenge. Pre-treating GBM with S100A4-TfR BsA may allow these promising treatments to persist and be more effective in mounting a strong ant-tumor immunity to tip the scale. While not knowing the precise mechanism of action for the BsA will not hinder clinical development, the precise mechanisms through which S100A4 inhibition reprograms the GBM TME will facilitate this effort. While others have reported that S100A4 binding to TLR4 or RAGE as important mechanisms(*77, 78*), we did not observe a significant difference in glioma growth or survival in *TLR4*-/- mice (unpublished), and there is very little RAGE expression on GBM TME cells to explain the therapeutic potential of S100A4 inhibition (unpublished). Nevertheless, consistent survival benefit offered by S100A4 inhibition in multiple ICI-resistant, clinically relevant models of GBM subtypes strongly suggest that S100A4 is a promising novel immunotherapy target for GBM.

### LIMITATIONS OF THE STUDY

This study provides robust pre-clinical evidence that targeting S100A4 genetically or therapeutically with an αS100A4/αTfR BsA in combination with ICI significantly improves overall survival in multiple mouse GBM models by remodeling the GBM TME and improving T cell infiltration via HEVs. While we find that BsA+αPD-1 may induce TLS-like structures in mouse GBMs, the precise mechanisms, including the S100A4 receptor and downstream signaling pathways mediating these changes in the TME remain incompletely defined and require further studies. Additionally, the underlying mechanism behind the heterogenous responses to genetic manipulation of *S100a4* and therapeutic treatment remain unresolved. Understanding the mechanism behind these differences is also important to help predict which patients may benefit from similar targeted therapy in a clinical setting.

## MATERIALS AND METHODS

### Mouse models and use

Mice were housed and handled in accordance with the protocols and procedures approved by the HMRI Institutional Animal Care and Usage Committees (protocol #IS00007185, #00001187 and #00001077). Mice were group-housed (up to five animals per cage) in a controlled environment with a standard 12-hour light/12-hour dark cycle. Animals had ad libitum access to food and water. We used male and female 8-12-week-old C57BL/6 mice or NSG (NOD.Cg-Prkdcscid Il2rgtm1Wjl/SzJ, JAX #005557) mice.

### Intracranial tumor injection and treatments

Starting 2 days prior to surgery and up to 2 days post-surgery, mice feed was switched to carprofen tablets. Immediately prior to surgery, buprenorphine SR (0.2mg/kg) were administered subcutaneously. Mice were then anesthetized via inhalation of 2.5% isoflurane and immobilized in a stereotactic apparatus. Freshly dissociated mouse glioma tumorspheres were injected into the mouse striatum using a stereotaxic device (Bregma: -0.5, 2.7, 3.5) as previously described. Animals were monitored daily and harvested once they began showing signs of neurological decline such as ataxia, severe lethargy, tilted heads, hunched posture. Depending on the experiment, mice were treated with 10 mg/kg αPD-1 (BioXCell: #BP0146), 10 mg/kg αCTLA-4 (BioXCell: #BP0131), 10 or 20 mg/kg αS100A4 (4-11) antibody(*57*), 10 or 20 mg/kg αS100A4/αTFR bispecific antibody, and equivalent concentration of Rabbit/Human chimaera IgG control or IgG/αTFR bispecific antibody control (produced in house). For mouse studies, exclusion criteria were severe complications during or immediately following intracranial surgery, such as wound dehiscence, severe ataxia, and infection. All procedures were approved by the HMRI Institutional Animal Care and Usage Committees.

### Primary mouse glioma tumorspheres lines

Primary mouse glioma tumorsphere lines were established from spontaneous GBMs and deeply characterized (Tran et al., in review). Specifically, we used *S100β-vERBb; p53^-/-^* SVP mouse model (lines 5459 and 6228), *tetO-EGFRviii; Ink4a/Arf^-/-^ ; Pten^-/-^ EIP* mouse model, *mGFAPcre; p53^flox/flox^; Pten^flox/flox^; PIK3CAH1047R^flox/+^*fPPP mouse model, *N/tv-a; PDGFA; shp53* KAB mouse model (KAB203), and *N/tv-a; PDGFA; shp53; shNF1* KABN mouse model (KAB200). Briefly, established glioma spheres from these models were intracranially injected into the striatum of C57BL6/J host mice. When the mice became symptomatic, they were euthanized and glioma regions were micro-dissected under a dissecting microscope. Single-cell suspensions were cultured in tumor stem cell medium consisting of DMEM/F12 (HyClone: #SH30023.FS) supplemented with 1X B27 (ThermoFisher: #17504044) and 1X GlutaMax (ThermoFisher: #35050061), or mouse neural stem cell (NSC) media consisting of TSC supplemented with 10 ng/ml mFGF (Peprotech: #450-33), and 20 ng/ml mEGF (Peprotech: #315-09). Mycoplasma testing was conducted periodically and prior to experimental use, including intracranial injections.

### Human GSC lines

Human GSC line MMC1 was previously reported (*39*) and cultured in human neural stem cell medium consisting of TSC supplemented with 10 ng/ml hFGF (Peprotech: #100-18C), and 20 ng/ml (Peprotech: hEGF #AF-100-15).

### Generation of sh-*Ctrl*/sh-*S100a4* knockdown cell lines

Lentivirus was produced in HEK293T cells maintained in DMEM (VWR: #16750-074) supplemented with 10% FBS. Cells were transfected with 9µg pCMV-ΔR8.91, 1µg pMD.G, and 10µg Hairpin-PLKO.1 in Opti-MEM supplemented with GlutaMAX (Thermofisher: #51985034) per 10 cm dish using lipofectamine 3000 (Thermofisher: #L3000008) according to the manufacturers protocol. Media was replaced after overnight incubation (approximately 16 hours), and viral supernatant was collected after 48 and 72 hours post-refreshment and stored at -80°C until use.

To generate short-hairpin(shRNA)-expressing cells, we utilized shRNA targeting scramble control l (sh-*Ctrl*) or *S100a4* (sh-*S100a4*), as previously described (*39*). Low passage primary mouse tumorspheres (<P10) were dissociated into single cells with Accutase (Corning: #25-058-CI) and spin-infected with shRNA lentivirus targeting scramble control (sh-*Ctrl*) or *S100a4* (sh-*S100a4*) prepared previously mixed with mNSC (1:1) at 1000 RCF for 1 hour at RT. After 16 hours of incubation, the media was replaced with fresh mNSC media containing puromycin (ThermoFisher: #A1113803)(500ng – 1µg/mL, depending on results from Puromycin-kill-curve) for selection. Puromycin-selected stable cell lines were maintained with 1 ug/mL puromycin until use. Successful knockdown of *S100a4* was validated via qPCR and western blot prior to use in experiments.

### Quantitative PCR

Total RNA was extracted using the TRIzol (Invitrogen: #15596026) phenol-chloroform phase-separation method and precipitated with isopropanol. cDNA conversion was done using the iScript™ gDNA Clear cDNA Synthesis Kit (Bio-Rad: #1725034) according to the manufacturer’s protocol. Quantitative PCR reactions were run in triplicate using the iTaq Universal SYBR Green Supermix (Bio-Rad: #1725120) according to the manufacturer’s protocol. Calculations were done by averaging triplicate quantification cycle (Cq) values, normalizing values to 18s RNA levels by subtracting the 18s Cq from the target gene’s Cq, and expression was calculated relative to the control.

### Construction and production of S100A4/TFR bispecific antibodies

S100A4-11 monoclonal antibody production and characterization was previously published(*57*). To generate the bispecific antibody, TfR scFv was fused to S100A4-11 using the same approach previously described(*52, 57*). In brief, the desired gene fragments were first amplified using PrimeStar GXL polymerase (Takara Bio: #R052A), and up to 3 fragments were then fused to create the whole or part of the novel antibody format using In-fusion HD cloning enzyme mix (Takara Bio: #102518). The heavy and light chain plasmids were co-transfected into Exp293 cells at equal weight ratios. Exp293-produced antibodies were purified using CaptivA Protein-A affinity resin (Repligen) and eluted with 0.1M glycine (pH=2.5) and then neutralized with 1/20 volume 1M Tris-HCl (pH=9). Buffer exchange to D-PBS was conducted with Amicon Ultra-15 ultrafiltration units (MW cutoff=30k) (MilliporeSigma: UFC901008D). We used Pierce High-Capacity Endotoxin Removal Spin Columns to remove Endotoxins (ThermoFisher: #88274) and measured endotoxin levels using an endotoxin detection kit (ThermoFisher: # A39552) and ensured all experiments are performed with preps with less than 1 EU/mg Ig.

### ELISA

For ELISA to assess brain and serum concentrations of the αS100A4/αTfR BsA or αS100A4 (4-11) mAb, high-binding plates (Corning: #9018) were coated with human S100A4 protein (Sino Biological: #10185-H08E) at 2µg/mL in PBS overnight at 4°C. Plates were then blocked in 40µm filtered 3% bovine serum albumin in PBS for 1 hour at room temperature. After blocking, plates were incubated in brain lysates (diluted 1:50), serum (diluted 1:20,000), and antibody standards for 2 hours at room temperature. All samples and standards were diluted in 1X ELISA diluent (Biolegend: #421203). After washing, plates were incubated with αHuman-IgG-HRP conjugate(1:5000) for 1 hour. 100 µLs of TMB substrate (ThermoFisher: #34022) were added for 10 minutes, and reactions were stopped with 2M H2SO4 solution (Biolegend: #423001). Plates were read at 450 nm absorbance.

### Cytokine array

Secreted cytokine and chemokine profiles were determined in the cell culture supernatants of the S100β-vERBb;p53-/- SVP mouse model (line 5459) using commercially available protein arrays (R&D Systems: #ARY028) according to the manufacturer’s instructions. We took conditioned media from sh-Ctrl or sh-S100a4 gliomaspheres cultured for 24 hours for analysis.

### siRNA transfection

20nM of siRNA (Universal Negative) were transfected into dissociated human gliomaspheres using Lipofectamine RNAiMAX Transfection Reagent (ThermoFisher: #13778075) according to manufacturer’s instructions. After 12 hours of transfection, the media was replaced with NSC media. After 48 hours, cells were used for downstream assays. Transfection was validated by using BLOCK-iT Fluorescent Oligo controls (ThermoFisher: #2013), and by assessing TfR protein expression by flow cytometry and by IF.

### Immune phenotyping by flow cytometry

Freshly dissected tumor tissues were micro-dissected into small chunks and then treated with a digestion buffer consisting of 1x Collagenase/Hyaluronidase (GeneDEPOT: #CA094-002) and DNAse I (GeneDEPOT: #D0100-050) diluted in Accutase for 20 minutes at 37°C. After 20 minutes, digestion buffer was removed, and the tissues were resuspended in TSC medium to triturate and generate single-cell suspensions. Cells were re-suspended in RBC lysis buffer to remove red blood cells. Following RBC lysis and washing with PBS, cells were strained through 40μm Flowmi cell strainers (VWR: #H13680-0040). Cells were blocked with TruStain FCX anti-mouse CD16/32 Antibody (Biolegend: #101320) then stained with multiple flow cytometry validated antibody cocktails (see below), and analyzed using either the BD Fortessa, LSRII, BD FACSymphony A5, or Cytek Aurora cytometers. Flow cytometry data was then analyzed and quantified using Flowjo (v10.10.1). All antibody dilutions and staining were performed in Brilliant Stain Buffer, and cells were incubated with a blocking solution containing mouse TruStain FcX (1:100) for 10 minutes prior to antibody staining.

### Immunofluorescence analysis

Thawed frozen sections or de-paraffinized sections underwent blocking in 5% goat or donkey serum (Sigma-Aldrich) with 0.2% TritonX-100 in PBS for one hour. After blocking, slides were incubated with primary antibodies in blocking buffer overnight at 4°C. Then, slides were incubated with appropriate Alexa Fluor secondary antibodies for 30 min at room temperature. Nuclei were counterstained with DAPI. The True VIEW autofluorescence quenching kit was used (Vector Laboratories: #SP-8400-15) to remove background fluorescence and to mount the slides. Images were obtained using a LEICA DM6 B Thunder upright microscope, an Olympus FV3000 confocal microscope, or a SLIDEVIEW VS200 universal whole slide imaging scanner. Immunofluorescence images were analyzed with custom pipelines in Qupath or CellProfiler. In brief, cells were segmented by DAPI signal using machine learning tools provided in Qupath or CellProfiler. For cell type identification, total signal per segmented cell was quantified, and cells were identified as positive or negative according to user-determined thresholds. For vessel area identification, pixels were identified as positive or negative according to user-determined thresholds and total positive pixel area was quantified. For region specific analysis, tumor regions were manually segmented based on nuclei density as determined by DAPI signal. Tumor border regions were then defined as projecting inwards and outwards by 100 µm from the tumor region outline – the remaining tumor regions were defined as tumor core. For IF experiments quantifying S100A4+ area, we used the MeasureImageIntensity module in CellProfiler to determine percent signal positive area based on a common user-defined threshold.

### COMET multiplexed IF imaging and analysis

Lunaphore COMET multiplexed immunofluorescence staining and imaging were performed at the University of Texas MD Anderson Flow Cytometry and Cellular Imaging Core Facility. De-paraffinized sections underwent heat-induced epitope retrieval using the Lunaphore HIER buffer according to manufactures instructions. Raw OME-TIFFs were imported into Visiopharm (v2025.08.1) without background subtraction. Image analysis including tissue region segmentation, cell segmentation, and positive-marker thresholding was performed using standard pipelines and tools in Visiopharm. Tumor regions were defined by Visiopharm machine learning pipelines trained on manually segmented tumor regions identified by nuclei density and Ki67+ staining. Tumor Border regions were then defined by expanding inwards and outwards 200 µm from the outlines of defined tumor regions. From these regions, individual cells were then identified via DAPI staining, and marker positivity was assessed using user-defined thresholds. Cell segmentation results were exported, to analyzed in, and visualized using R.

### GLASS dataset analysis

Clinical data and bulk RNA sequencing was downloaded from the Glioma Longitudinal Analysis consortium via Synapse (syn17038081; release date 2025-09-30). We used the accompanying metadata for annotation of IDH status and transcriptional subtype. Transcriptional subtype was assigned to each sample aliquot as the signature with the lowest ssGSEA p-value.

For survival analysis, each patient contributed one tumor sample. *S100A4* expression was log2(TPM+1) transformed. For all glioma, IDH-wildtype, and IDH-mutant cohorts, the earliest profile sample for each patient was used. For the primary diagnosis cohorts, the primary tumor was used. For the recurrent cohort, the earliest profiled recurrence sample was used. Patients were divided into *S100A4*-high and -low cohorts by median within each cohort. Endpoints for overall survival from initial diagnosis was used for all gliomas and IDH-mutant gliomas; endpoint for overall survival from profiled surgery was used for IDH-wildtype gliomas. Survival was compared using Kaplan-Meir curves with the log-rank Mantel cox test.

For S100A4 expression analysis, S100A4 expression was compared between IDH-wildtype, IDH-mutant 1p19q non-codeletion, and IDH-mutant 1p19q co-deletion samples. Primary and recurrent samples were compared within patients using the paired Wilcoxon signed-rank test. For comparison between transcriptional subtypes, we used the unpaired Wilcoxon rank-sum test.

Association between S100A4 expression status and survival was estimated with multivariable Cox proportional hazards regression. Covariates were applied to the model if recorded for at least 70% of patients, had more than one category, and had at least 5 patients in each category. Models were fitted on complete cases. Age was presented per 10-year increase. Hazards ratios with 95% confidence intervals and Wald *p*-values were reported.

Multivariable binary logistic regression was performed in each cohort to assess clinical features associated with S100A4 expression status. Predictors were the same covariates used within each cohort’s Cox model. Regression coefficients (log-odds) with Wald 95% confidence intervals were reported.

Analysis was performed in R (v4.4.1), survminer (v0.5.2), survival (v3.8-3), data.table (v1.16.4), and ggplot2 (v3.5.2).

### Bulk RNA-seq analysis

Bulk RNA sequencing analysis was conducted with an in-house pipeline as described previously(*79*). Gene-level counts were imported, and Ensembl identifiers were mapped from symbols using org.Mm.eg.db. Samples were assigned to various treatment arms and analyzed with edgeR (v4).

### Gene ontology analysis

For each sample, gene log2CPM values were scored against mouse MSigDB collections retrieved from msigdbr using gene set variation analysis (GSVA) with a Gaussian kernel and gene set sizes limited to 10-500 genes (MSigDB modules H, C2, C5, C8). Enrichment scores matrices were fitted with limma, and gene sets passing BH-adjusted p < 0.05 were retained for heatmap display.

Over-representation analysis (ORA) was performed by taking up and down regulated genes (FDR < 0.05, |log2FC| > 1) and testing against all expressed genes as background. All tests used BH adjustment with p < 0.05 and q < 0.2.

GO enrichment network plot was determined using differentially expressed genes between tested groups (FDR < 0.05 & |log2FC| > 1). Differentially expressed genes were split into up- and down-regulated lists, tested separately for GO_BP enrichment against all expressed genes as background (Benjamini–Hochberg p < 0.05). Each node represents a GO term, and node area is proportional to number of differentially expressed genes annotated to the term. Each pie plot shows proportion contributed by up- and down-regulated lists. Terms sharing gene membership at Cohen’s κ > 0.3 are connected by an edge, with edge width proportional to κ; terms with less than two connections were omitted. Clusters were delineated by random-walk clustering, labelled by their most significantly enriched member term.

### Cell type deconvolution from bulk RNA-seq

Enrichment scores for 64 cell types were estimated with xCell (v1.1.0). Enrichment scores were mean centered across samples for display.

### Proteomics analysis

Label-free proteomic abundances were exported as a protein-by-sample matrix of relative abundances. Redundant protein groups from shared peptides were collapsed before testing if gene symbols shared a common prefix and values were identical across all samples. Proteins were retained if detected above the floor in at least two samples of at least one group. Floor values were set to missing and abundances were log2 transformed. Missing values were imputed per sample by drawing from a normal distribution downshifted into the low tail of that sample’s observed distribution under a fixed random seed. Differential abundance was assessed with limma. Variance was moderated by empirical Bayes with an intensity-dependent prior trend and robust hyperparameter estimation. Proteins with BH-adjusted p < 0.05 were considered differentially abundant. Gene set variation analysis was performed as described for bulk RNA-seq data.

### Visium spatial RNA-seq sequencing

Formalin-fixed paraffin embedded (FFPE) samples were sectioned and taken to determine the DV200 value using Qiagen’s RNeasy FFPE Kit value of the sample according to the manufacturer’s instructions (Qiagen: #73504). After the tissues were confirmed to be of acceptable quality with a DV200 value >30%, one slide per sample was used for H&E stain and to generate special transcriptomics libraries using the Visium CytAssist, Visium CytAssist Slide and Cassette 6.5mm kits, Visium FFPE Reagent Kits, and Visium Mouse Transcriptome Probe Kits according to the manufacturer’s instructions (10X Genomics, Pleasanton, CA). All FFPE sectioned samples and required reagents were loaded onto the Visium CytAssist for probe capture and extension, followed by amplification, adaptor, and index ligation according to the manufacturer’s instructions. Barcoded spatial transcriptome libraries were sequenced on Novogene’s NovaSeq X Plus platforms. Raw sequencing reads were aligned to the mm10 genome (mouse) using Space ranger from 10X genomics with default parameters.

### Visium spatial RNA-seq analysis

Filtered feature-barcode matrixes and paired H&E images were imported into R with Seurat. Spots with less than 200 detected UMIs were removed as quality control. Retained spots were normalized with SCTransform. Select gene expression was plotted using the GroupHeatmap function from SCP package showing mean row-z-scores.

Cell-type deconvolution analysis of Visium samples was performed using the Spotiphy package (v0.3.1)(*80*) in Python (v3.9.23). We read visium data for each sample into an AnnData object using scanpy (v1.10.3). Spot-level annotations generated in Seurat were imported from an exported metadata table and mapped back onto the AnnData object. We imported a single cell RNA-seq reference we generated from our analyzed and annotated single cell RNA-seq dataset of intratumor cells, as described in the previous section “Single cell RNA-seq analysis”. For cell-type deconvolution, we used the scRNA-seq reference to estimate the per-spot proportion of each cell type by variational inference and visualized per-spot proportion results as a spatial feature plot. Circle colors that denote proportions were normalized within each sample to that sample’s 90th percentile and capped at 1.

### Copy number alteration (CNA) analysis

We inferred copy-number alteration (CNA) profiles for each Visium spot across eight samples using the R package infercna (v1.0.0), sorting genes by chromosomal location and calculating a CNA score using a 150-gene sliding window, adapting from methods published described previously(*81*). For our normal karyotype reference, we combined two independent normal mouse brain Visium datasets publicly available from 10x Genomics. .

### Tissue region assignments for Visium samples

To classify tissue region assignments for each spot, we assigned “malignant” spots as classified by CNA analysis as “Tumor”, “non-malignant” spots as “Normal Brain”, and remaining spots as “Other”. We then fit a 2D kernel density estimate to the TumorCore spots (MASS::kde2d, bandwidth scaled 0.7×). After testing multiple parameters, we defined the contour enclosing 92% of the spots taken as the tumor outline; interior rings were dropped so each outline was filled completely, and multiple outlines were retained where a section contained more than one focus. All spots inside an outline were labelled as TumorCore. We then defined zones on the hexagonal array. The outmost layer of spots outlining each tumor region was expanded inward and outward 2 layers to define the TumorBorder. All remaining spots were classified as NormalBrain.

### Non-negative Matrix Factorization analysis

De novo transcriptional programs were identified with GeneNMF (v0.9.2). Non-negative matrix factorization was run per sample across ranks k = 4–12. Per-sample programs were consolidated into consensus metaprograms by clustering programs by cosine similarity of their gene weights and retaining enough genes per program to explain 50% of the weight. Metaprograms encompassing less than 10 genes were dropped. Metaprogram genes were tested for enrichment against MSigDB collections (MSigDB modules H, C2, C5, C8) using all detected genes as universe. We assigned functional and biological labels to metaprograms based on their constituent genes and enriched gene sets. We used metaprogram genes to calculate signature scores for each metaprogram for each spot with the sigScores() function from the scalop package (v1.1.0). Signature scores were determined as the mean expression of signature genes after gene-wise centering across all spots, thus reflecting relative expression and being comparable across genes of differing baseline abundance. We dropped signatures for which fewer than 50% of member genes were detected in the matrix. We assigned each spot to a metaprogram based on the top signature score calculated for that spot. We then assessed CNA malignancy bins proportions by assigned metaprogram to determine whether metaprograms were classified as malignant or non-malignant.

For neighborhood analysis, Vasc-high spots were first identified by taking all spots, ordering by Vasc MP signature scores determined previously, and setting a threshold for “Vasc-high” spots at two standard deviations above the mean. We then extracted coordinates for these spots, and for each spot determined the surrounding six spots as the neighborhood. The neighborhood score for a given gene set or module was calculated as the mean signature score for all spots in the neighborhood.

### Statistical analysis

Statistical comparison was using Graphpad Prism (v11.0.2) (GraphPad Software, La Jolla) or R (v4.4.1). Values and error bars represent mean ± standard error mean (SEM). Respective number of replicates are indicated in the figures or figure legends. *p*-values were determined by an appropriate statistical test such as Student’s t-test or analysis of variance (ANOVA) and indicated in the figure legends.

## Data and code availability

Sequencing data generated in this study well be deposited to GEO upon publication. This article also analyzes existing, publicly available data, and the sources for these datasets are listed in the materials and methods section. This article does not report original code. Any additional information required to reanalyze the data reported in this article is available from the lead contact upon request.

## ACKNOWLEDGEMENTS

We thank Dr. Sonia Villapol for providing access to the VS200 slide scanner at Houston Methodist Research Institute. We also thank the staff of the Comparative Medicine Program, Methodist Flow Cytometry Core, and Advanced Cellular and Tissue Microscopy Core at Houston Methodist Research Institute, and the Flow Cytometry and Cellular Imaging Facility at University of Texas MD Anderson for their assistance and support. We also thank Dr. Xuejun Fan for supporting the ELISA assay development for the BsA, and all members of the Yun lab for support and helpful suggestions.

## AUTHOR CONTRIBUTIONS

KY: Conceived the study, designed experiments, analyzed data, and prepared the article.

TW, JL, MF: Designed experiments, collected and analyzed the data, and prepared the article.

NA, SJY: Designed experiments, and collected and analyzed the data.

WX, FL, YA, DJ, HT, SG, RK: Collected and analyzed the data.

ZZ, NZ: Provided reagents, designed experiments, and reviewed the article.

---

**Supplementary Figure 1:**
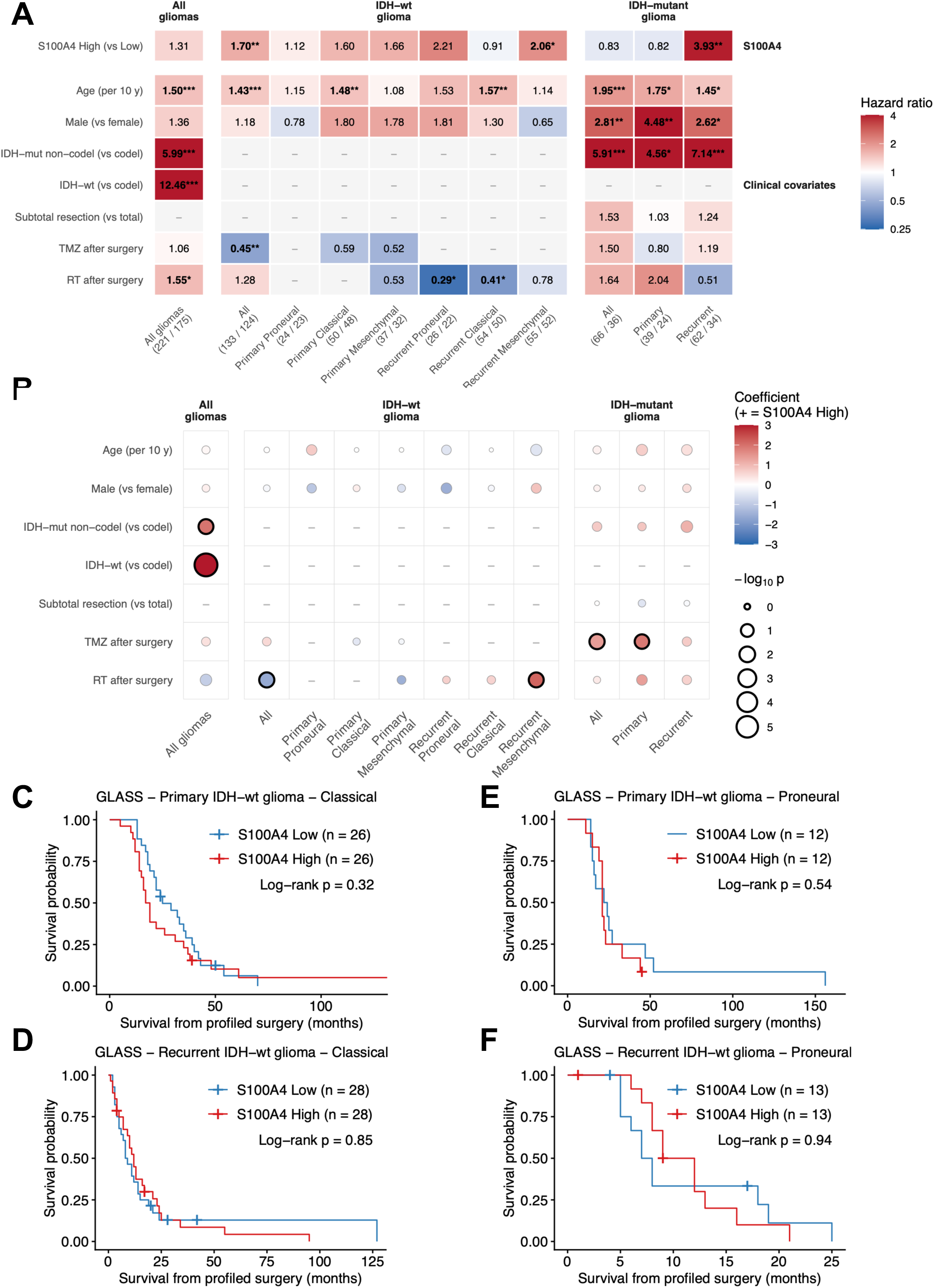
S100A4 expression in human glioma. **A.** Heatmap depicting results of multivariable cox proportional hazards models of overall survival in GLASS glioma cohorts. Rows represents clinical variables and columns represent separate models/cohorts. Numbers below each column indicate number of patients present in KM curve and deaths at last follow up. Color and number in each cell indicates hazard ratio for each clinical variable adjusted for all other variables present in the column. Age is per 10 years. IDH-wildtype cohorts were analyzed from the date of the profiled surgery, all others from diagnosis. *p < 0.05, **p < 0.01, ***p < 0.001 (Wald test, unadjusted for multiple comparisons). **B.** Dotplot depicting results of multivariable logistic regression models for S100A4 high and low expression. Rows represents clinical variables and columns represent separate models/cohorts. Color indicates the regression coefficient (log-odds), and dot size indicates - log10p, black outlines mark p < 0.05. Age is per 10 years, reference categories shown in parenthesis. **C-F.** Kaplan-Meier plot depicting overall survival distributions from profiled surgery of (C) primary and (D) recurrent IDH-WT Glioma Classical subtype and (E) primary and (F) recurrent IDH-WT Glioma Proneural subtype, stratified by high or low S100A4 expression, split determined by median value within cohort. *p*-values determined by log-rank Mantel-cox test.

**Supplementary Figure 2:**
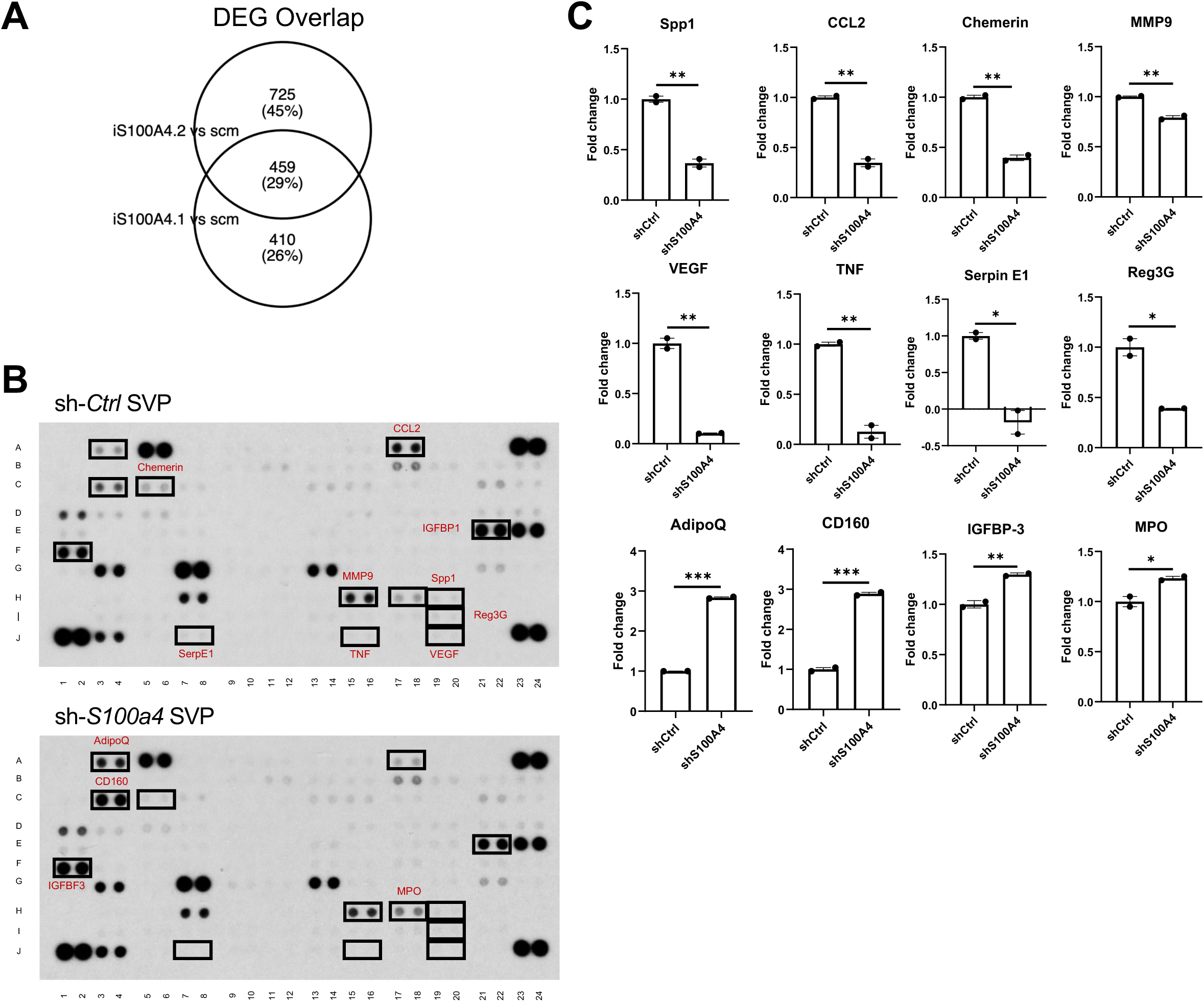
*S100A4* knockdown regulates cytokine expression in human GBM cells and modulates the GBM TME. **A.** Overlapping differentially expressed genes from bulk RNAseq data of human MMC1 gliomaspheres treated with control or two siRNAs (siS100A4.1 and siS100A4.2 ) targeting *S100A4*. **B, C.** Secreted cytokine and chemokines in conditioned medium collected from sh-*Ctrl* and sh-*S100a4* SVP (5459) cells (24 hours) determined by mouse XL cytokine array kit (R&D system). 8 and 4 soluble proteins were separately reduced and induced by S100a4 inhibition. Downregulated proteins included Spp1, Ccl2, Chemerin, MMP9, VEGF, TNF, Serpin E1, and Reg3G. Upregulated proteins include AdipoQ, CD160, IGFBP-3, and MPO.

**Supplementary Figure 3:**
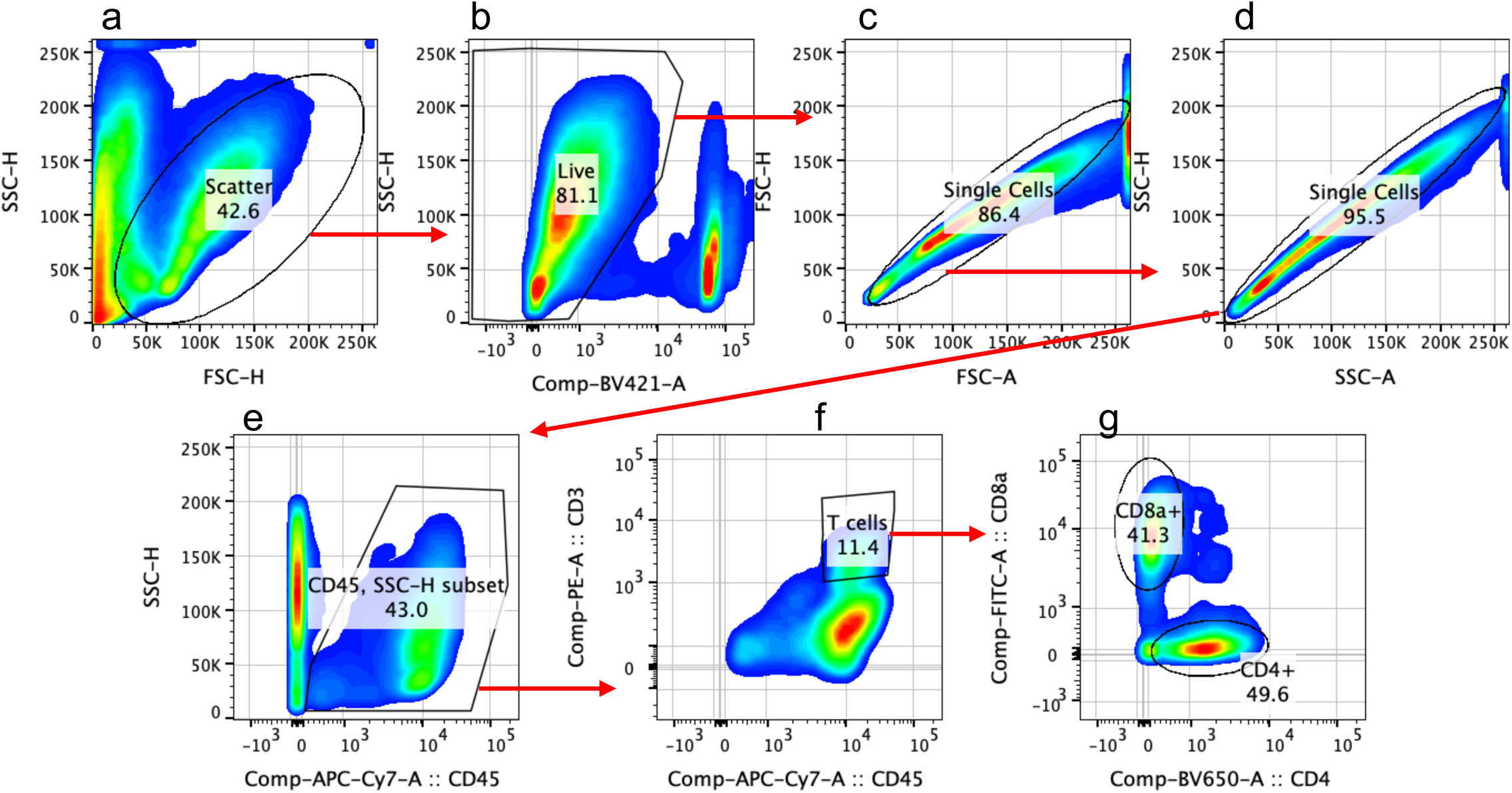
Representative flow cytometry gating for all T cells, CD4 T cells, and CD8 T cells. Flow cytometry gates for quantification of all T cells, CD4 T cells, and CD8 T Cells from 5459 tumors. Flow cytometry gates were manually determined in Flowjo and applied to all samples. T cells were selected via sequential gating by exclusion of cellular debris (a), dead cells (b), doublets (c, d), CD45-cells (e), and CD45-CD3-cells (f). CD4 and CD8 T cells were selected based on differential non-overlapping expression of CD4 and CD8.

**Supplementary Figure 4:**
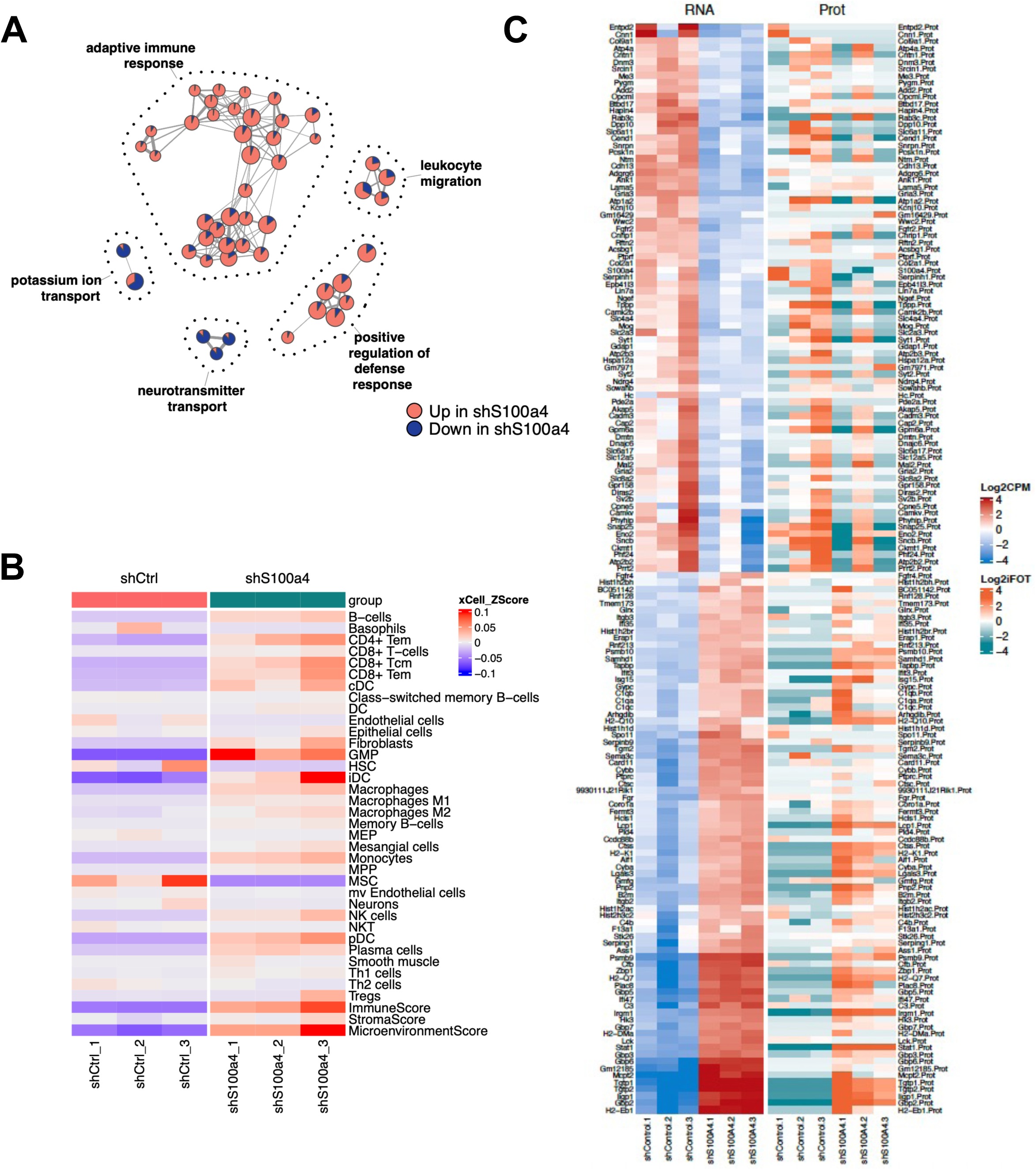
Analysis of sh-*Ctrl* and sh-*S100a4* SVP samples. **A.** GO enrichment network plot of differentially expressed genes between sh-*Ctrl* and sh-*S100a4* SVP (5459) tumors. Each node is a term, represented by a pie chart that shows proportion contributed by the up- and down-regulated genes, with area representing number of differentially expressed genes annotated to the term. Dotted outlines delineate communities identified by random-walk clustering, labelled by most significantly enriched member term. **B.** Heatmap displaying xCell deconvolution of immune and stromal cell type abundance in sh-*Ctrl* and sh-*S100a4* 5459 tumors at endpoint. Color intensity reflects Z scores. **C.** Heatmap displaying concordant gene and protein level expression from bulk RNA sequencing and mass spectrometry, respectively, of sh-*Ctrl* and sh-*S100a4* SVP samples.

**Supplementary Figure 5:**
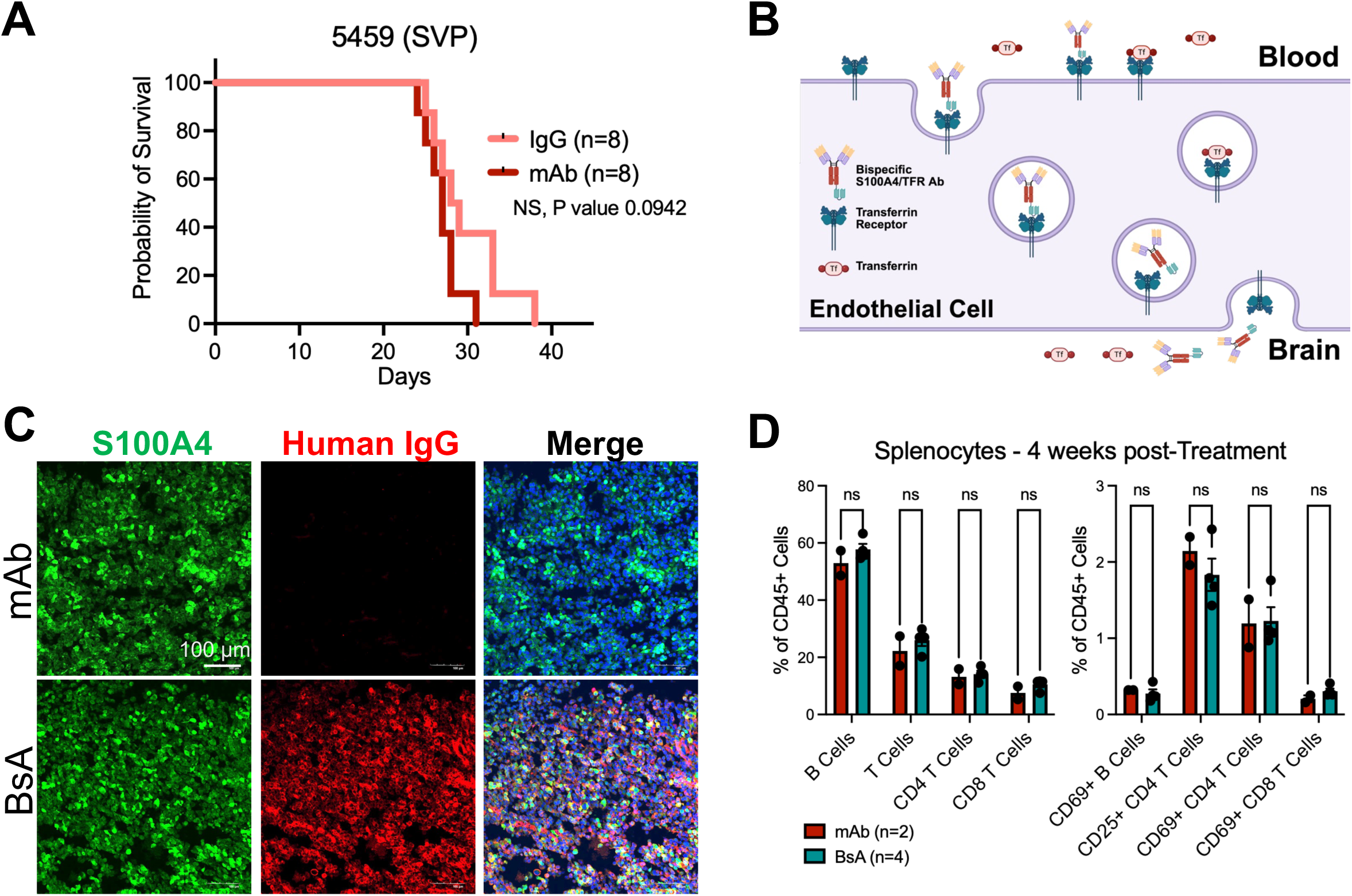
BsA S100A4/TFR schematic and treatment in mouse GBM. **A.** Kaplan-Meier survival curve of B6 mice intracranially injected with 5459 gliomaspheres and treated with mAb or IgG control. *p*-value determined by logrank Mantel-Cox test. (n=8/group). **B.** Schematic of BsA transferrin receptor mediated endocytosis across the blood brain barrier. **C.** Representative IF confocal images of mouse 5459 glioma stained for S100A4 and Human IgG at 24 hours post treatment with S100A4 monoclonal or BsA antibody. BsA has human Fc region and can be detected with anti-human Fc. **D.** Flow cytometry analysis of splenocytes 4-weeks from B6 mice after one-time mAb (n=2) and BsA (n=4) treatment. *p*-values determined by multiple unpaired t-tests.

**Supplementary Figure 6:**
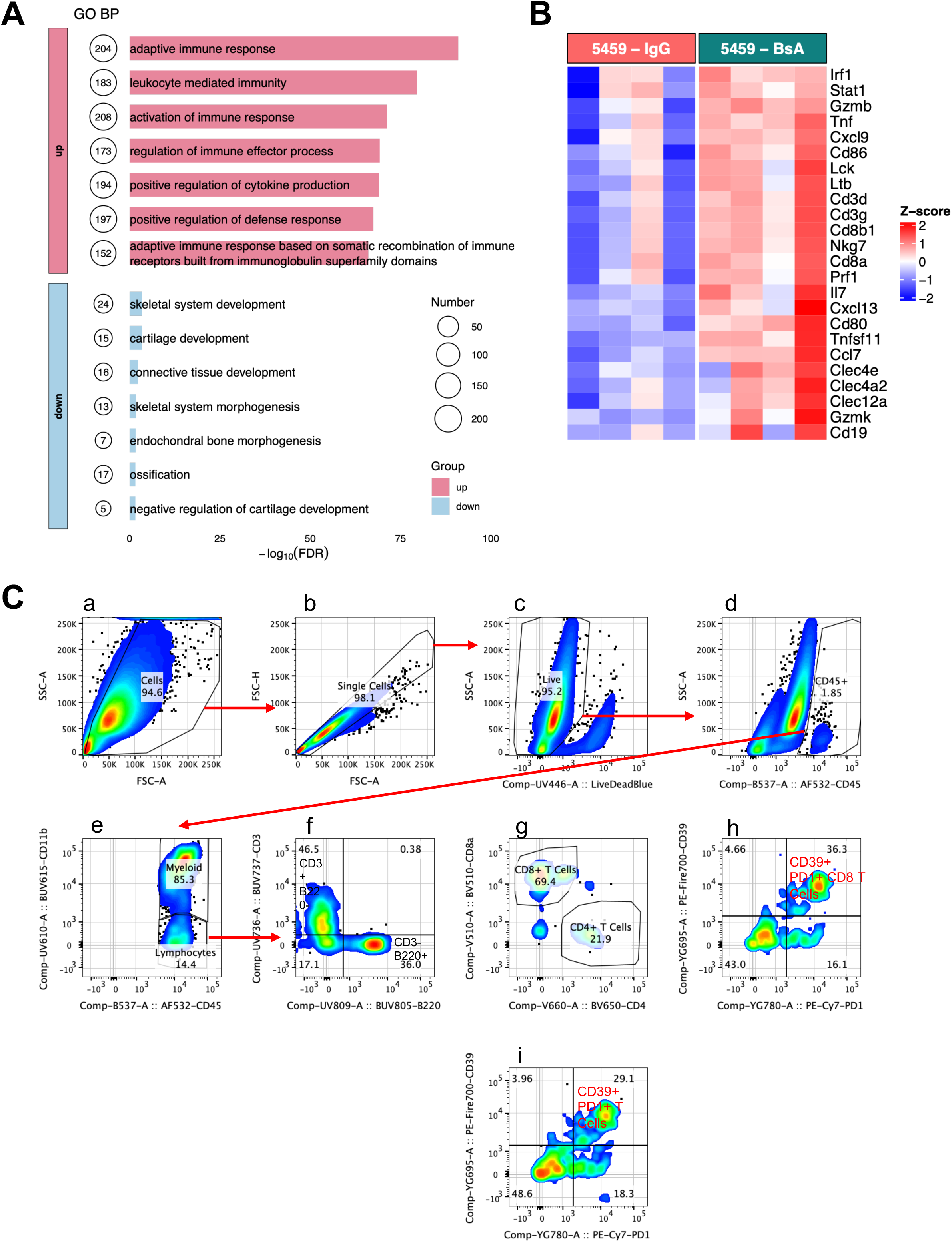
BsA treatment in mouse GBM remodels the GBM TME. **A.** Over-enrichment analysis showing up and down regulated GO_BP gene sets for BsA treated 5459 tumors vs controls. **B.** Heatmap of selected up and downregulated genes associated with innate and adaptive immune response in IgG and BsA treated 5459 tumors. **C.** Flow cytometry gates for quantification of CD39+ PD1+ T cells and CD8 T cells. Flow cytometry gates were manually determined in Flowjo and applied to all samples. T cells were selected via sequential gating by exclusion of cellular debris (a), doublets (b), live cells (c), CD45-cells (d), and CD45+CD11b-cells (e). T cells were selected based on expression of CD3 and lack of B220 (f), and CD8 T cells were selected based on expression of CD8 and lack of CD4. Tumor reactive T cells were determined from T cells (i) and CD8 T cells (h) based on expression of CD39 and PD1.

**Supplementary Figure 7:**
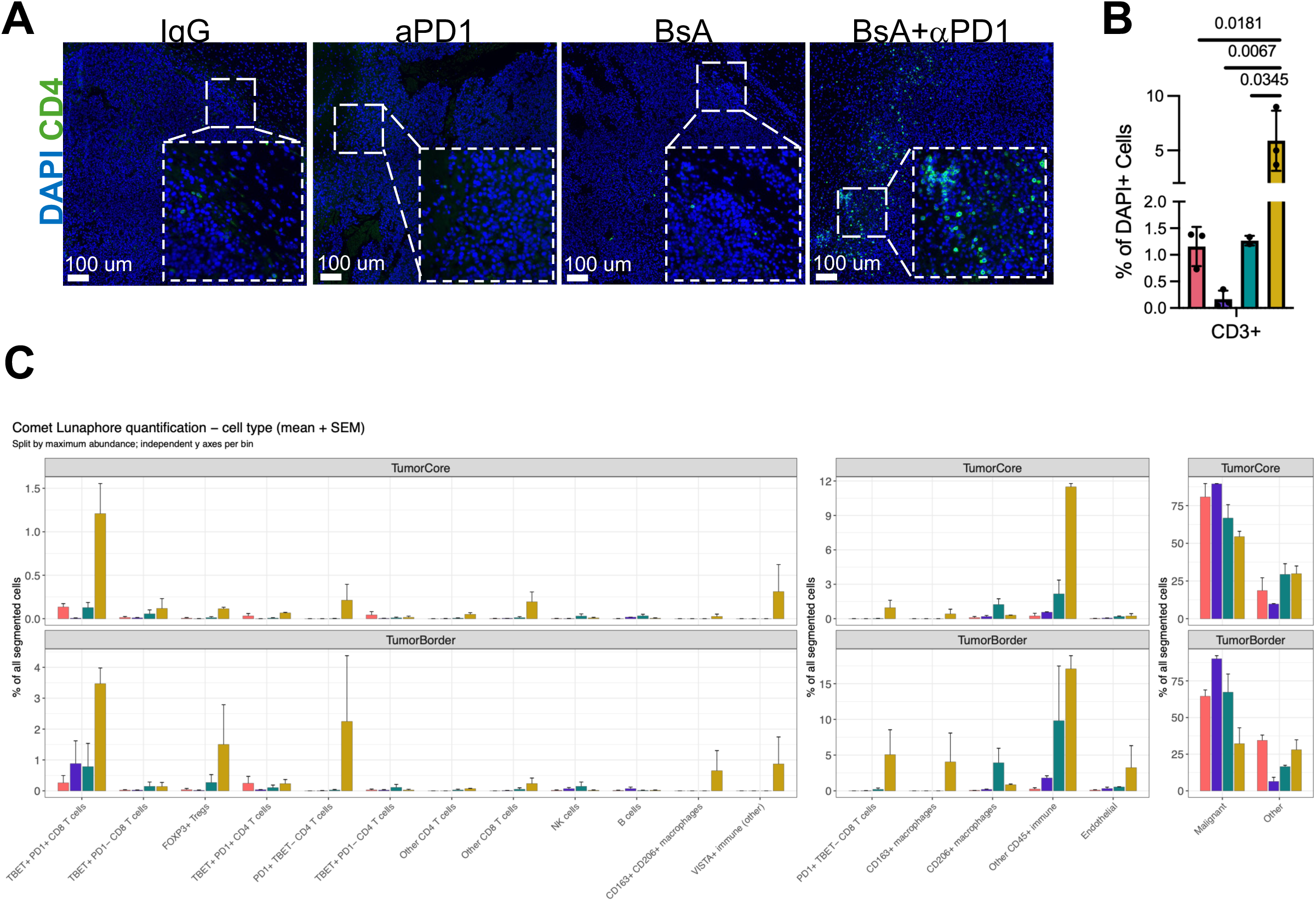
BsA+αPD-1 treatment increases T cell infiltration. **A.** Representative IF images for CD4 T cells from IgG, BsA and Combo treated endpoint 5459 samples. Scale bars represent 100 µm. **B**. Quantification of IF analysis from endpoint 5459 tumors. n=3-4 per group. *p*-values determined by one-way ANOVA with Šidák correction for multiple comparisons, p < 0.05 displayed. Bar plots and error bar indicate mean and SEM, respectively. **C.** Bar plots comparing cell type abundance as a percentage of total segmented cells, grouped by treatment group and split by tissue region. n=2/group.

**Supplementary Figure 8:**
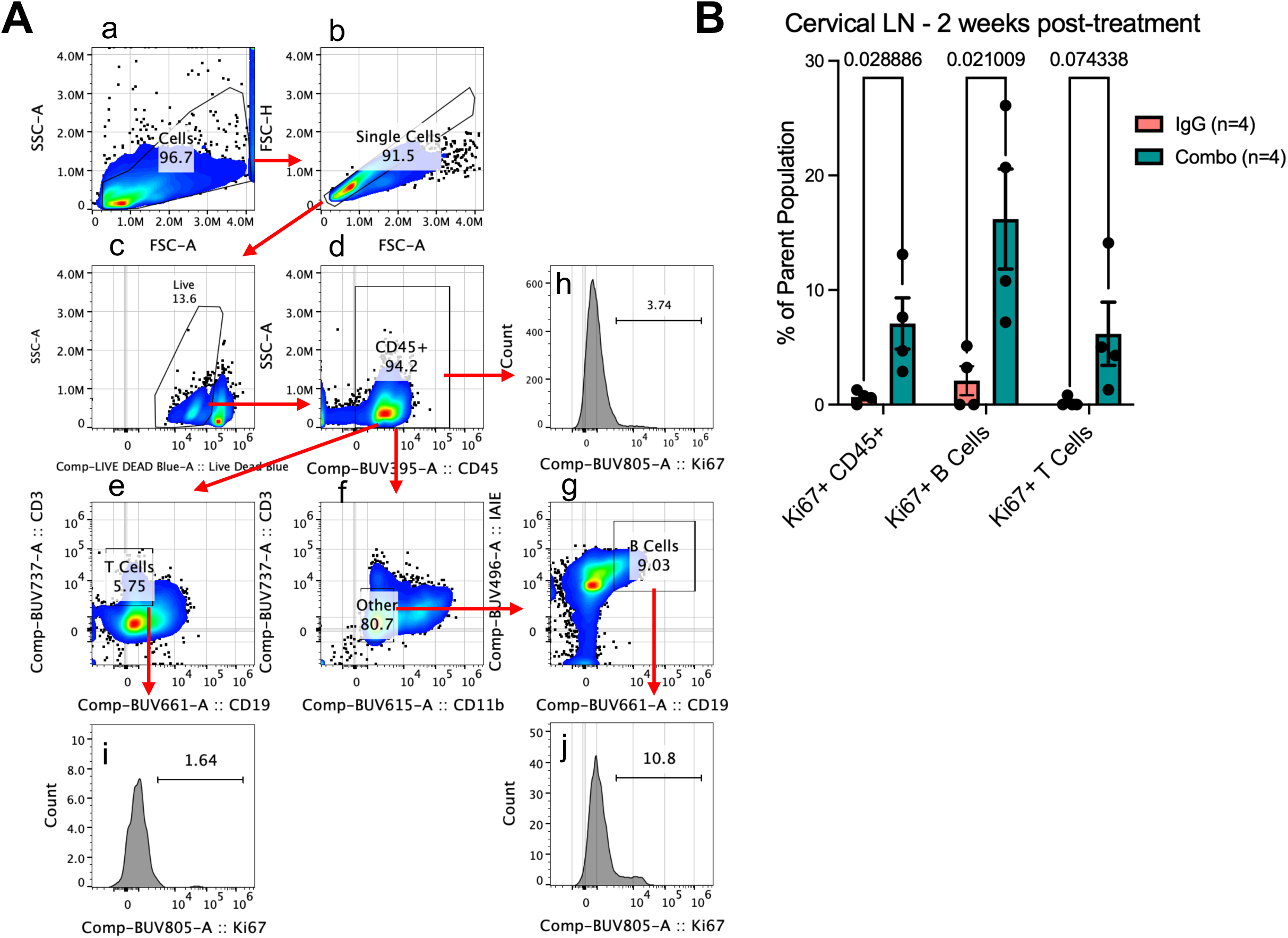
BsA+αPD-1 treatment activates peripheral immunity. **A.** Flow cytometry gates for quantification of Ki67+ cells. CD45+ cells were selected via sequential gating by exclusion of cellular debris (a), doublets (b), live cells (c), followed by selection of CD45+ cells (d). T cells were further selected by expression of CD3 and lack of CD19 (e). B cells were selected by exclusion of CD3+ and CD11b+ cells from CD45+ cells (f), followed be selection of CD19+ IAIE+ cells (g). Ki67+ gates were selected via histogram for CD45+ cells (h), T cells (i), and B cells (j). **B.** Flow cytometry analysis of immunes cells in cervical lymph nodes from IgG or Combo treated 5459 tumor bearing mice at 2 weeks post-treatment. Values are displayed as percentage of parent population, e.g. Ki67+ CD45 cells displayed as a percentage of total CD45+ cells, all cells are gated off LiveDeadBlue-CD45+ populations, LiveDeadBlue-CD45+ CD11b-CD3-CD19+ IAIE+ populations, and LiveDeadBlue-CD45+ CD19-CD3+ populations, n=4/group. *p*-values determined by unpaired two tailed t-test.

**Supplementary Figure 9:**
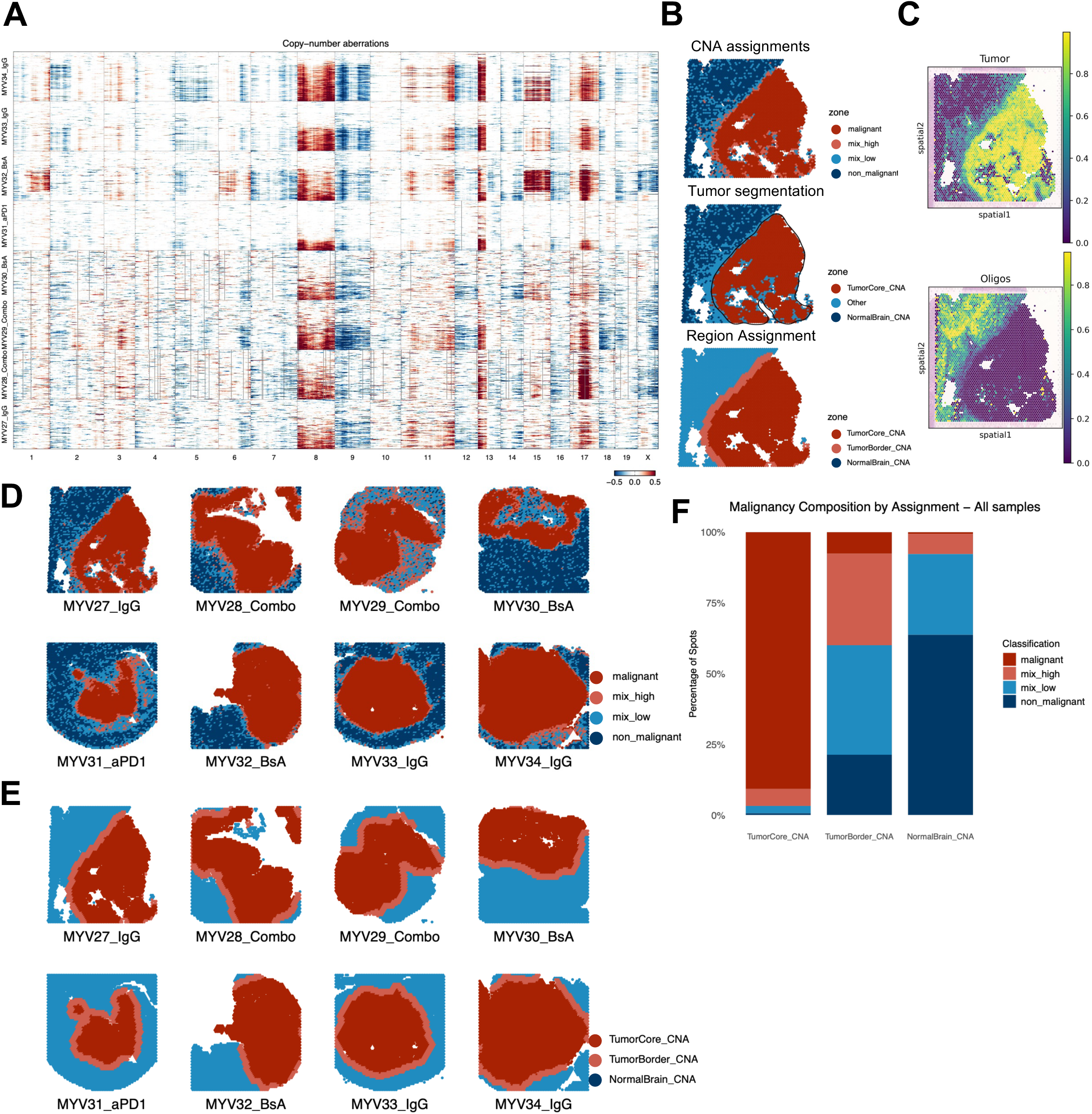
Copy number aberration analysis of BsA treated mouse GBM tissues. **A.** Copy number aberration (CNA) plot showing relative copy-number profile inferred from Visium spot expression. Each row represents a spot, each column is a gene ordered by genomic coordinate, segmented by chromosome region. Color indicates inferred relative copy number, with red indicating copy number gain and blue indicating copy number loss. Spots are grouped by sample. **B.** Representative spatial dimension plot showing spot level assignment of CNA malignancy bins, tumor tissue segmentation assignments, and final tissue region assignments. **C.** Representative spatial feature plots showing spot level cell level deconvolution of glioma tumor and oligodendrocyte proportions. **D, E.** Spatial dimension plot showing spot level assignment of CNA malignancy bins (D) and tissue region assignments (E) for all samples. **F.** Stacked bar plot showing proportions spots assigned to each malignancy bin in each tissue region.

**Supplementary Figure 10:**
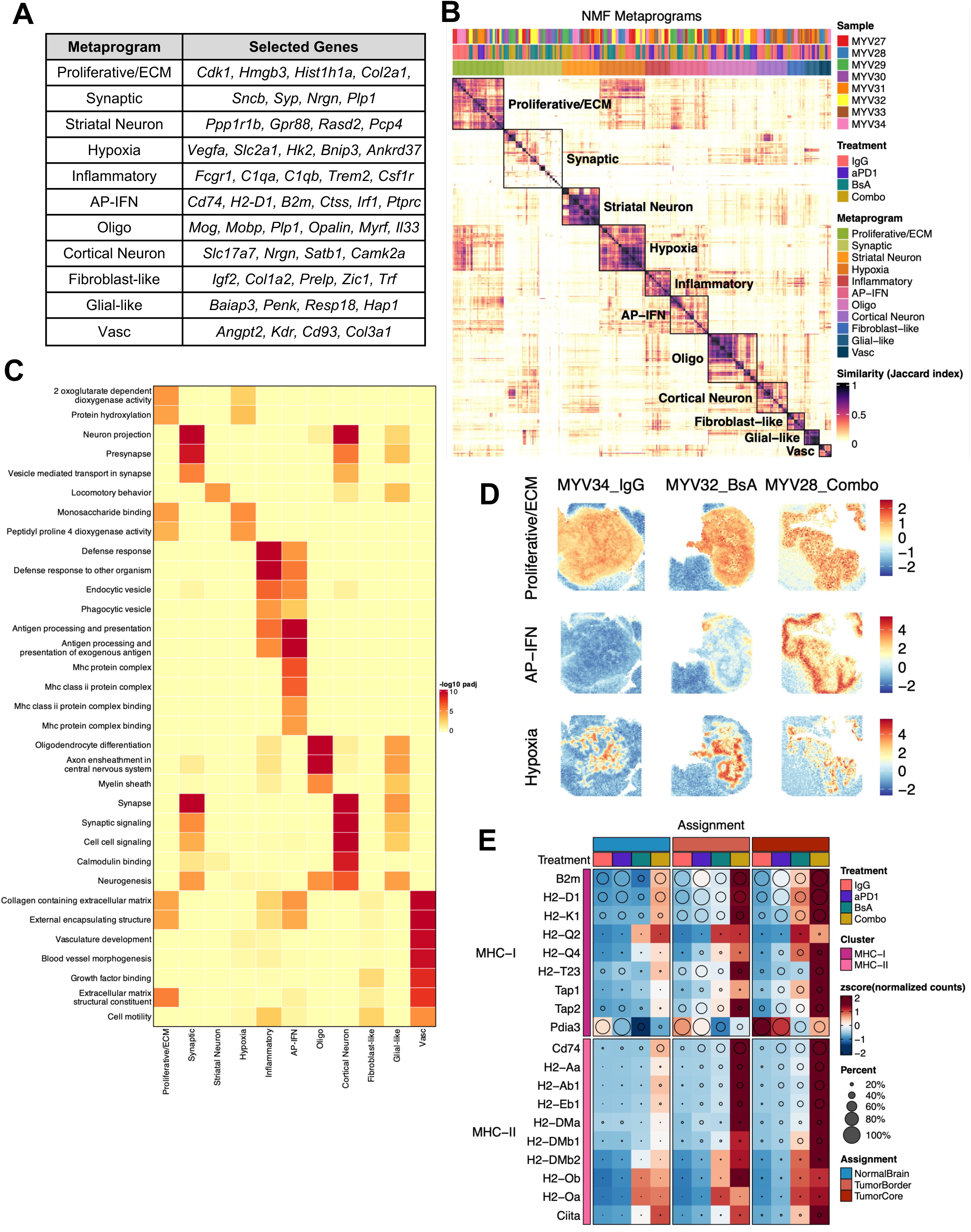
nNMF analysis of BsA treated mouse GBM tissues. **A.** Table showing nNMF MPs and signature genes. **B.** Symmetric heatmap of pairwise Jaccard similarity between individual Visium non-negative matrix factorization (nNMF) programs derived from each sample. Rows and columns represent individual NMF programs, color indicates Jaccard index measuring overlap between genes of each pair. Identified and annotated consensus metaprograms (MP) are outlined and shown. **C.** Heatmap showing results from over representation analysis (ORA) of consensus nNMF MP genes for GO_BP, GO_CC, and GO_MF pathway gene sets. **D.** Spatial feature plots showing spot level Proliferative/ECM, AP-IFN, and Hypoxia MP scores for three samples from IgG, BsA, and Combo treatment groups. Color for each spot reflects the metaprogram score, as indicated by the color scale. **E.** GroupHeatmap showing gene expression of MHC-I and MHC-II antigen presentation associated genes, grouped by treatment group and split by tissue region.

**Supplementary Figure 11:**
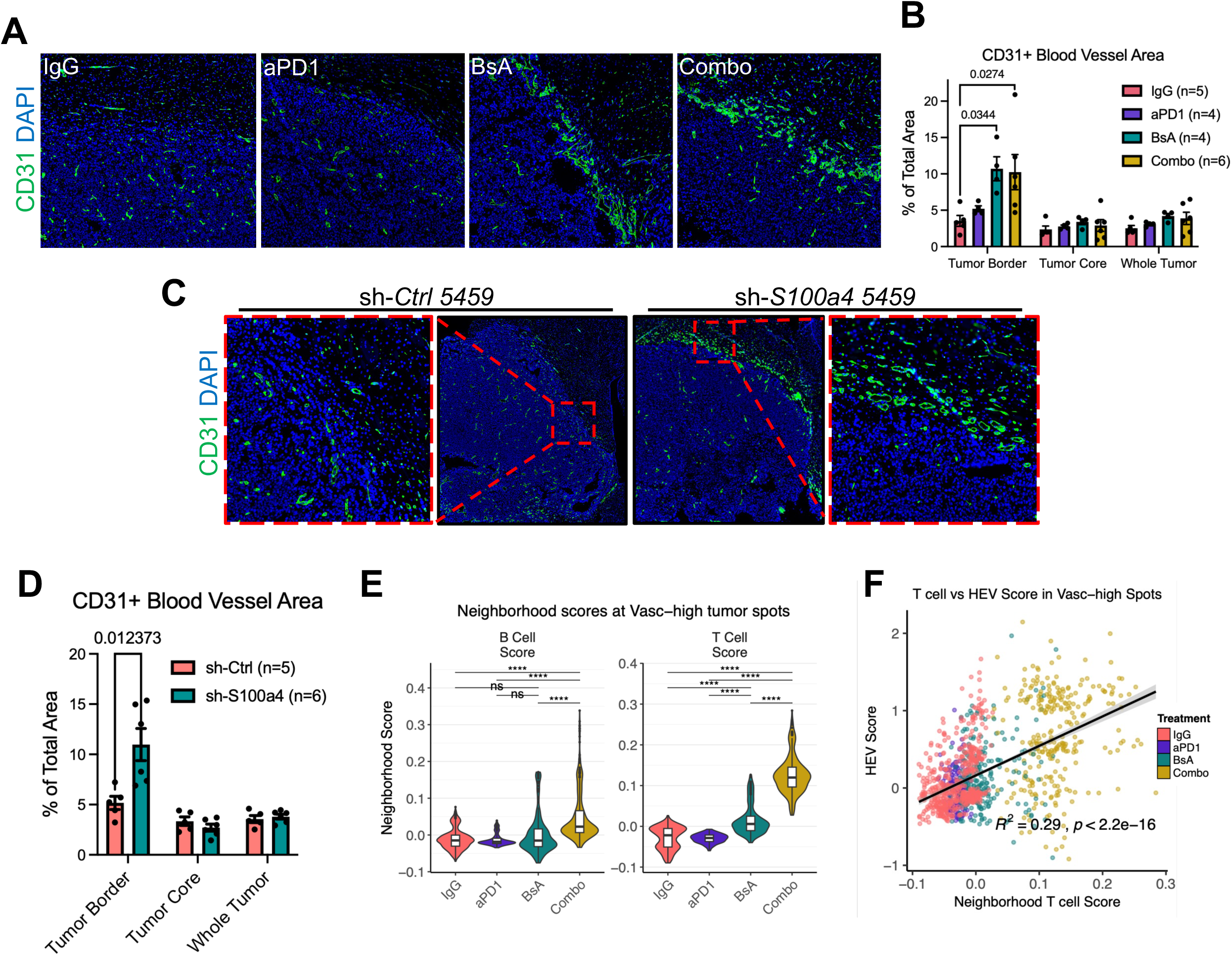
Vasculature analysis of *S100a4*-KD and BsA treated mouse GBM tissues. **A.** Representative IF microscopy images of endpoint IgG, αPD-1, BsA, and BsA+αPD-1 treated 5459 tumors stained for CD31. **B.** Quantification of CD31+ area as a percentage of total area, by tissue regions. *p*-values determined by one-way ANOVA with Šidák correction for multiple comparisons, p < 0.05 displayed. Bar plots and error bar indicate mean and SEM, respectively. **C.** Representative IF microscopy images of endpoint sh-*Ctrl* and sh-*S100a4* 5459 tumors stained for CD31. **D.** Quantification of CD31+ area as a percentage of total area, by tissue regions. *p*-values determined by unpaired two-tailed t-tests, p < 0.05 displayed. Bar plots and error bar indicate mean and SEM, respectively (n=5-6 per group). **E.** Violin plot showing neighborhood scores of B and T cell gene modules in Vasc-high expressing spots from the tumor border and tumor core, grouped by treatment group. Box represents median and interquartile range. Vasc-high spots were determined taking all tumor border and tumor core spots from all samples, computing a signature score from Vasc nNMF MP genes, and thresholding at 2 standard deviations above the average. **F.** Pearson correlation analysis of T cell and HEV neighborhood scores in Vasc-high spots.

